# Pol γ possesses separate metal binding sites for polymerase and strand displacement functions

**DOI:** 10.64898/2026.01.25.701366

**Authors:** Noe Baruch-Torres, Joon Park, Josue Mora-Garduño, Arkanil Roy, Anupam Singh, G. Andrés Cisneros, Luis G. Brieba, Smita S. Patel, Y. Whitney Yin

## Abstract

Accurate replication of mitochondrial genome (mtDNA) integrity, which is essential for cellular metabolism and energy supply, relies primarily on DNA polymerase gamma (Pol γ), Twinkle helicase, and mitochondrial single-stranded DNA binding protein (mtSSB). Twinkle alone exhibits little helicase activity while reports indicate that Pol γ displays from modest to limited unwinding activity. This led us to dissect Pol γ strand displacement activity using structural, biochemical and in silico approaches. Here, we show that human Pol γ carries out robust strand displacement synthesis at physiological concentrations of divalent metal ions which reveals that distinct metal-binding sites can independently regulate DNA synthesis and unwinding activities. We further showed that Pol γ can displace RNA/DNA hybrid with comparable efficiency as DNA/DNA duplex, representing a key implication on RNA primer removal to preserve mtDNA integrity. Our cryo-electron microscopy structures of Pol γ complexed with a template containing downstream dsDNA and an incoming nucleotide revealed the structural mechanism for the strand displacement activity. We identified four conformational states that represent successive stages of DNA unwinding, accompanied by coordinated rearrangement of the downstream DNA and Pol γ elements that mediate strand displacement. This work establishes biochemical and structural mechanisms of Pol γ strand displacement activity, providing fundamental insight into human mitochondrial DNA replication and integrity.

**Graphical abstract:** 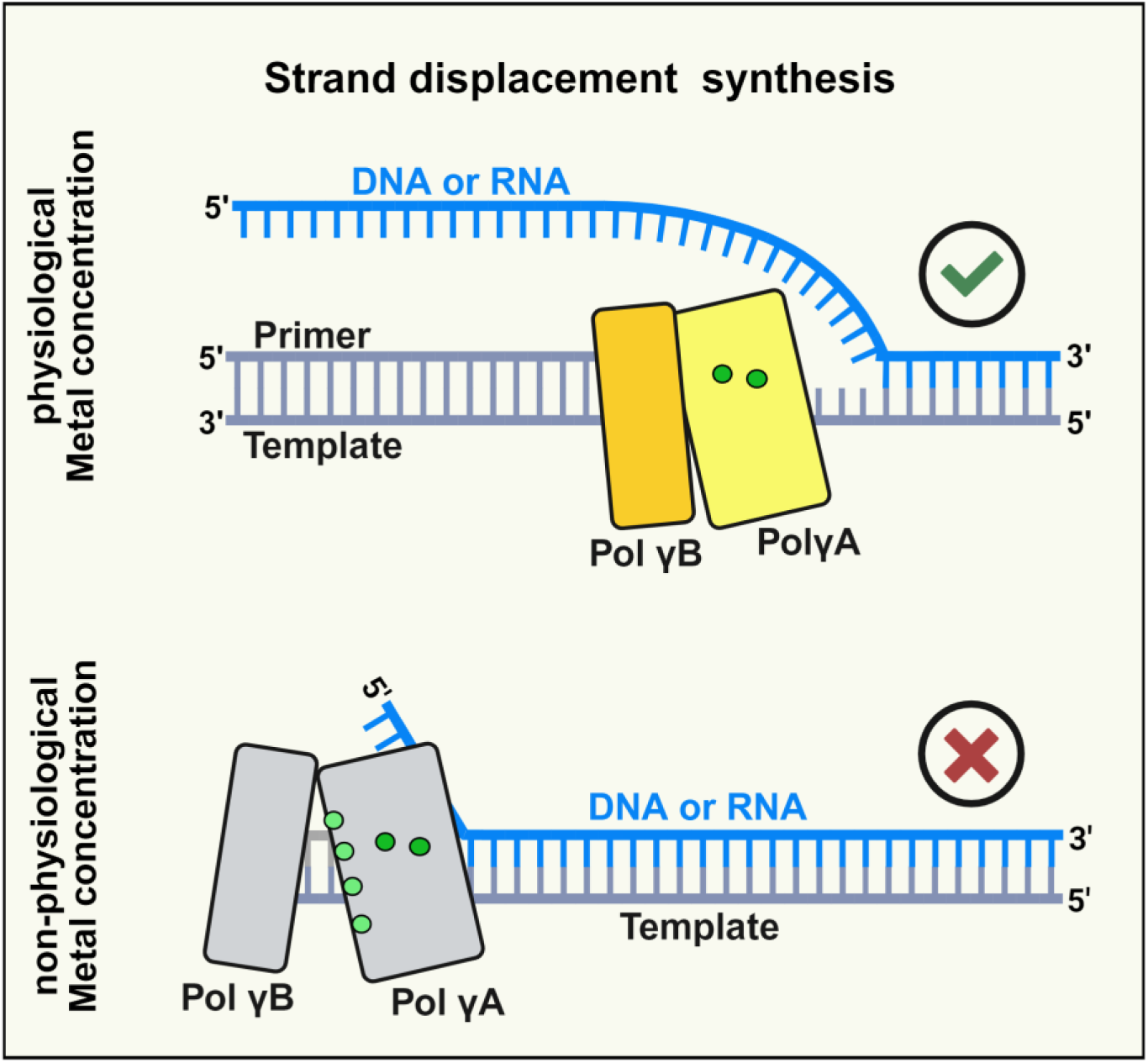

## INTRODUCTION

Mitochondrial DNA (mtDNA) integrity is essential for cellular metabolism and energy supply. Human mtDNA is a 16.9 kbp double-stranded circular DNA, whose two strands (H- and L-strands) are replicated primarily via an asymmetrical, continuous mechanism (1–3), and based on the biological context, strand-coupled/RITOLS mechanism (4–6). Replication of double-stranded human mtDNA is accomplished by a core replisome consisting of Twinkle helicase, DNA polymerase gamma (Pol γ), and single-stranded DNA binding proteins (mtSSBs). Mutations in Pol γ and Twinkle catalytic core have been implicated in neurological and muscular disorders, as well as aging (7–9), illustrating their importance in cellular and organ functions.

A classical division of labors in replisome is that the helicase unwinds dsDNA powered by the energy from ATP hydrolysis while DNA polymerase synthesizes using the newly generated single-stranded DNA template. Nevertheless, human Twinkle helicase alone exhibits poor DNA unwinding activity, and in fact, it effectively anneals DNA (10, 11). Pol γ and Twinkle jointly conduct dsDNA synthesis and unwinding efficiently (12), raising the possibility that Pol γ may act as a driving molecular motor for the replisome by converting chemical energy from dNTP incorporation to unwind DNA duplex, an activity termed strand displacement synthesis.

Lacking a designated organellar primase, the RNA primers that initiate human mtDNA replication are synthesized by the sole mitochondrial RNA polymerase (POLRMT) (13–15). The RNA primers synthesized by POLRMT are longer than the typical lengths, and their removal is critical for mitochondrial genome integrity (16, 17). A critical step for RNA primer removal involves Pol γ strand displacement activity to generate flaps which are then cleaved by nucleases to maintain mtDNA intact (18–20, 21). Previous investigations have demonstrated that Pol γ exhibits from moderate to reduced strand displacement synthesis activity. An *in vitro* biochemical study has shown that Pol γ performs limited strand displacement activity to create short 5’-flaps in downstream DNA and RNA primers, although RNA displacing is more inefficient (<10 nt) in comparison to DNA (16, 17). While using a mini circle DNA containing a preformed fork DNA demonstrated that Pol γ is unable to continue synthesis on dsDNA region (22). A report using single-molecule optical tweezer combined with biochemical approaches describes a robust strand displacement synthesis at the first base pair of the DNA but subsequently reduces by ⁓30-folds as Pol γ is unable to prevent the non-template strand reannealing (23). Under both approaches Pol γ exo+ strand displacement processivity resulted similar (37 ± 15 nt). In contrast, Pol γ exo-deficient variant has been reported to have strand displacement activity allowing to continue synthesis on dsDNA substrates (24), indicating the polymerase possesses intrinsic elements for molecular motor function that is modulated by the exonuclease activity (24, 25). In this work we aimed to elucidate the biochemical and structural basis governing Pol γ strand displacement activity.

We report here that the metal ions that support Pol γ catalysis can also regulate its dsDNA unwinding activity. We also propose that Pol γ contains at least two distinct types of metal binding sites: the high-affinity sites supporting Pol γ enzymatic activity and DNA unwinding, whereas the low-affinity sites suppress DNA unwinding without affecting DNA synthesis activity. Thus, under physiological concentrations of Mg^2+^ or Mn^2+^, Pol γ displays robust strand displacement synthesis with high fidelity and with processivity comparable to that on a single stranded template. We also demonstrate that Pol γ can displace RNA primers with high efficiency comparable to DNA primer removal and it is supported by physiological concentrations of either Mg^2+^ or Mn^2+^. We solved cryo-EM structures of Pol γ in complex to dsDNA, revealing three protein elements that would be involved in strand displacement activity: the helix-turn-helix catcher domain, the splitter helix and the strand displacement helix. Altogether, our study sheds light on the factors regulating Pol γ strand displacement activity and its implication in DNA replication and RNA primer removal to preserve mtDNA integrity.

## Materials and Methods

### Expression and purification of recombinant proteins

His-tagged Pol γA and Pol γB were expressed in *Sf9* insect cells and *E. coli* Rosetta (DE3) cells, respectively, and purified as previously described (21). Briefly, both Pol γA wild-type and exonuclease deficient variants were purified to homogeneity using TALON (Cytiva, Marlborough, MA) and gel filtration column Superdex 200 columns. Pol γB was purified via a Ni-NTA (Qiagen, Germantown, MD), Mono S chromatography. Purified PolγA and Pol γB were mixed at 1:2 ratio and purified on a Superdex 200 column. Fractions containing Pol γ holoenzyme were pooled, concentrated and aliquoted for storage at −80°C.

T7 DNAP exo+ was purchased from New England Biolabs (catalog number M0274S).

### Nucleic acid substrates

The oligonucleotides were purchased from Integrated DNA Technologies, Inc (Coralville, Iowa). A 5’-FAM-labeled 30-nt primer (ATT GGA AGT AGG GAT AGT CCC GAA CCT CGC) were annealed to a 74-nt template (5’-Biotin AGT GCT TAC ACC TGC CGC ATG ATC ACG GTA CGA GCT TGC TTT AGG CGA GGT TCG GGA CTA TCC CTA CTT CCA A/Inverted dT-3’) with or without the 85-nt non-template (5’-TTT TTT TTT TTT TTT TTT TTT TTT TTT TTT TTT TTT TTT TTT TTT AGC AAG CTC GTA CCG TGA TCA TGC GGC AGG TGT AAG CAC/Inverted dT-3’) to generate a p/t or a fork DNA.

The gap-constructs containing DNA blocker or RNA blocker were formed by annealing a 40-nt DNA blocker (5’-AGC AAG CTC GTA CCG TGA TCA TGC GGC AGG TGT AAG CAC T-3’) or a 40-nt RNA blocker (5’-AGC AAG CUC GUA CCG UGA UCA UGC GGC AGG UGU AAG CACU-3’) to the 5’-FAM-labeled 30-nt primer/74 template.

Circular 3.2 kb ssDNA was produced as reported previously with minor changes (26). Briefly, 5 mL of 2x YT media supplemented with tetracycline/ampicillin was inoculated with XL1-Blue strain carrying pGEM3Zf(+)-HOM vector and incubated at 37°C for 12 h to increase the cell culture density. The culture was diluted with fresh 2x YT media containing VCSM13 helper bacteriophage (Agilent, Santa Clara, CA) at 6×10^7^ pfu/mL and incubated at 37°C for 1 h. To select the infected cells, kanamycin was added, and the incubation continued for another 18 h. The phage particles were precipitated and utilized for ssDNA extraction using E.Z.N.A.® M13 DNA Mini Kit (Omega BIO-TEK), following manufacturer instructions.

The *midi circle* DNA substrate was formed by annealing the 5’-cy5 71-nt primer to the 3.2 kb circular ssDNA at 75°C for 5 min then decrease the temperature to 22°C at a rate of 1.0°C per 90 s in a Bio-rad T100 thermocycler (Hercules, CA).

### Strand displacement assay on short fork DNA substrates

200 nM Pol γ exo+ or exo– variant was mixed with 100 nM DNA substrate in a buffer GB (20 mM Hepes pH 8.0, 105 mM NaCl, 35 mM KCl, 2.5 mM DTT, 4% glycerol, 0.1 mgmL^−1^ BSA), DNA synthesis was initiated by addition of 0.2 mM dNTPs and Mg^2+^ or Mn^2+^ at indicated concentrations. The reaction mixture was incubated at 37°C for 10 min, then quenched by adding Q buffer (80% Formamide, 50 mM EDTA pH 8.0, 0.1% SDS, 5% glycerol, and 0.02% bromophenol blue). T7 DNAP exo+ was assayed similarly except in reaction buffer containing 20 mM Tris-Cl pH 7.5, 50 mM KCl, 1 mM DTT and 200 nM of enzyme. The samples quenched were heated at 95°C for 5 min, then placed on ice and applied to a 17% polyacrylamide gel and 8M urea electrophoresis. The gel was imaged on a GE Typhoon 950 scanner and quantified using ImageQuant software TL (GE Healthcare). The percentage of full-length products was calculated by the intensity ratio of the full-length product to the sum of the rest.

### Mismatch proofreading assays

5’-^32^P-labeled 26-nt primer (MM-p26 primer: 5’- AT ATT ATT TAC ATT GGC AGA TTC AAT-3’) was hybridized to a 40-nt MM-Temp40 template (5’-AA TCT AGT CCC <u>AAG CTT</u> GAA TCT GCC AAT GTA AAT AAT AT-3’) to generate a DNA substrate containing a T/C mismatch at the 3’-OH end of the extending primer. As controls, a 40bp DNA containing a T/C mismatch were formed by annealing a 40-nt oligo (5’-^32^P-AT ATT ATT TAC ATT GGC AGA TTC AA**<u>T</u>** CTT GGG ACT AGA TT-3’) to the MM-Temp40 template, and a 40bp Watson-Crick complementary DNA duplex was formed by annealing a 40-nt oligo 5’-^32^P-AT ATT ATT TAC ATT GGC AGA TTC AA**<u>G</u>** CTT GGG ACT AGA TT-3’) to the MM-Temp40 template. Only the 40bp W-C dsDNA contains a Hind III site.

500 nM of the MM-p26/ MM-Temp40 was preincubated with 100 nM of Pol γ exo+ in GB buffer (20 mM Hepes pH 8.0, 105 mM NaCl, 35 mM KCl, 2.5 mM DTT, 4% glycerol, 0.1 mgmL^−1^ BSA) at 37°C for 5 min. The reaction was started by addition of 0.2 mM dNTPs and 0.32, 0.64, 1.25, 2.5 or 5 mM MgCl2 and quenched after 20 min incubation by adding 1:10 volume of Q buffer. A duplicated set of reactions were stopped by boiling at 95°C for 5 min and cooled down gradually to 20°C followed by the Hind *III* addition to monitor the Pol γ exo+ mismatch excision. Samples were boiled at 95°C for 5 min and resolved on a TBE 17% polyacrylamide denaturing gel. DNA bands were quantified using ImageQuant software TL (GE Healthcare) and plotted as the percentage of cleaved products at each Mg^2+^ concentration.

### Tread-milling analyses

Pol γ dNTP incorporation and excision from the 3’-end of the primer were measured on a 70-nt minicircle substrate annealed to a 110-nucleotide primer as described previously (27, 28). The DNA substrate was incubated with Pol γ on ice for 30 minutes, then at 37°C for 5 minutes. Reactions were initiated by addition of 50 µM dGTP, 200 µM each of dATP, dTTP and dCTP, trace amount of α-^32^P-dGTP in the presence of 50 mM Tris-HCl pH 7.5, 40 mM NaCl, 10% glycerol and 2 mM DTT and from 0.1 to 5.0 mM MgCl2. All reactions were quenched after 10 minutes with addition of 4 M formic acid. 1 µl of each of the quenched reactions were spotted on polyethyleneimine-cellulose TLC sheet (Millipore). Analytes were resolved on the TLC with potassium phosphate buffer (pH 3.8). Dried TLC sheets were used to expose phosphor screens overnight. Screens were scanned on Typhoon FLA 9500 scanner and spot intensities corresponding to dGMP incorporated in the newly synthesized DNA, unused dGTP, dGDP and excised dGMP were quantified using ImageQuant TL program (Cytiva).

### Unwinding RNA/DNA hybrid

A gapped construct mimicking when replication reaches an RNA primer was composed by annealing the 74-nt template to the 5’-FAM-labeled 30-nt primer and a 40-nt RNA blocker oligo. We run primer extension assays containing 200 nM Pol γ exo+ and 100 nM of RNA or DNA blocker substrate in a buffer GB. Reactions started by the addition of 0.2 mM dNTPs and Mg^2+^ or Mn^2+^ at indicated concentrations. Reactions were stopped after 10 min incubation at 37°C by adding 9 folds of Q buffer (80% Formamide, 50 mM EDTA pH 8.0, 0.1% SDS, 5% glycerol, and 0.02% bromophenol blue). Samples were resolved on 17% polyacrylamide and 8M urea gel. The gel was imaged on a GE Typhoon 950 scanner and quantified using ImageQuant software TL (GE Healthcare). The percentage of full-length products was calculated by the intensity ratio of the full-length product to the sum of the rest.

Using optimal magnesium concentration for strand displacement activity (0.64 mM), we repeated the strand displacement assays and quenched from 15 to 600 s. Samples were analyzed on a 17% polyacrylamide denaturing gel. The full-length extension bands were measured using ImageQuant software TL (GE Healthcare) and plotted as function of time.

### In silico simulation of Polγ divalent metal binding sites

The pol γ system was modeled using the 4ZTZ crystal structure (29), focusing on the catalytic subunit Pol γA. The missing regions were filled with Rosetta Fold, generating multiple structures. The best model was selected based on visual inspection and RMSD alignment with the crystal structure as previously described (30).

Parameters for the incoming nucleotide were taken from prior work (31). Protonation states were set with ProPKA (32), at pH 7, and structural checks and hydrogen addition were done using MolProbity (33). The system was neutralized and solvated in AMBER18 with ff14SB (34), OL15 (35, 36), and TIP3P force fields. Based on system volume, 72 Mg²⁺ ions and 144 Cl⁻ ions were added to achieve 20 mM MgCl₂, whereas three Mg²⁺ ions and 2 Cl⁻ ions were in the 1 mM system. Simulations were carried out for each of five created 1 mM systems to reduce bias.

MD simulations were run in AMBER18 using pmemd.cuda (37). Minimization involved 10, 000 cycles (steepest descent and conjugate gradient). Heating to 300 K was done with Langevin dynamics (38), using a 2 ps⁻¹ collision frequency and initial restraints of 100 kcal mol⁻¹Å⁻², then gradually reduced. The systems with 20 mM concentrations had a production run of 500 ns in triplicates (for a total of 1.5 μs), and the 1 mM concentrations had a production of 100 ns, in quintuplets, due to the random placement of the ions (for a total of 500 ns). All bonds involving hydrogen were treated with SHAKE (37). Long-range Coulomb interactions were handled with smooth Particle Mesh Ewald (PME) method (39) and long-range van der Waals interactions are approximated using default isotropic correction in AMBER (39) with a 10 Å cutoff. Parameters used for calculations and starting co-ordinates were present in zenodo (https://zenodo.org/records/4469899). Analysis of the simulations was done using the CPPTRAJ suite of AMBER.

### Competition metal binding assays

200 nM Pol γ was mixed with 100 nM DNA and incubated for 5 min at 37°C, then 0.2 mM dNTPs and varied Ca^2+^ (0 to 10 mM) with or without constant concentration 0.32 mM Mn^2+^ (or 0.64 mM Mg^2+^) were added and incubated for 10 min. The reaction mixtures were quenched with Q buffer at 1:10 ratio. Reaction products were resolved on a TBE 17% polyacrylamide gel containing 8M urea. The gel was imaged on a GE Typhoon 950 scanner, and products were analyzed using the ImageQuant software TL (GE Healthcare).

### Midi-circle template replication assays

Assays were carried out with 7 nM of the 3.2 kbp *midi-circle* DNA substrate and 35 nM Pol γ in a GB buffer. The mixture was incubated for 10 min on ice and 5 min at 37°C before adding 0.2 mM dNTPs and 0 to 10 mM MgCl2. Reaction mixtures were incubated for 15 min, quenched by addition of 6X QB solution (18% Ficoll 400, 6% SDS and 120 mM EDTA pH 8.0), and subjected to electrophoresis on a 0.8% alkaline gel at 20V for 17 hours.

The gels were soaked in AB buffer (1M Tris-Cl pH 7.6, 1.5 M NaCl) for 45 min in rocking motion. To stain the circular ssDNA template and ladders, neutralized gels were incubated with SYBR gold dye (1x final concentration) for 60 min in 1X TAE buffer. Gels were scanned in a GE Typhoon 950 imaging system using dual-wavelength mode at 635 nm to visualize the Cy5 fluorophore on the primer and at 488 nm to visualize SYBR gold-stained DNAs.

### Cryo-EM Sample Preparation

A 77-nt dumbbell DNA (5’P-GCT TTT CTG GTG AAA AGC TGG TCG GCA GCG CTT GAG CAG CGG CAG CTG GTG CTG CCG CTG CTC AAG CGC TGC CGA C/ddC/-3’) was synthesized with 5’ phosphate and 3’ dideoxycytosine and purified with high performance liquid chromatography by Integrated DNA Technologies. It was annealed by heating 95°C for 5 minutes and slowly cooled overnight.

To prepare the complex for cryo-EM studies, 2 µM of Pol γ holoenzyme formed by Pol γA exo– and Pol γB ΔI4 (21) was incubated on ice with an equimolar amount of 77-nt dumbbell DNA in buffer containing 25 mM HEPES pH 7.5, 140 mM KCl, 10 mM CaCl2, 10 mM βME, 0.01% Octyl-β-Glucoside, and 1 mM dATP. The sample (4 µL) was applied to a plasma-cleaned QUANTIFOIL R 2/1 Cu 200 grid and rapidly frozen in liquid ethane using a Vitrobot Mark IV system at 22 °C and 100% humidity (Thermo Fisher Scientific).

### Cryo-EM Data Acquisition and processing

The frozen grids were loaded into a Titan Krios G3i (Thermo Fisher Scientific) equipped with K3 direct electron detector with BioQuantum energy filter (15-eV energy slit) (Gatan) and operated at 300 keV at Stanford SLAC Cryo-EM Center. Cryo-EM data were automatically acquired using EPU software in counted super-resolution mode at a nominal magnification of ×105, 000 (corresponds to 0.43 Å/pix) with a nominal defocus range between −1.5 and to −2.5 μm. Forty-frame movie stacks were collected over 2-s exposure with a total dose of 49.60 e^−^/Å. A total of 10, 227 movie data were collected.

The movie frames were imported into cryoSPARC (40) for image processing (Fig. S3). Movies were motion-corrected and 2× binned using Patch Motion Correction, resulting in 0.86 Å/pix. Contrast transfer function (CTF) of resulting micrographs was determined using patch CTF estimation. After discarding micrographs with CTF fit higher than 4 Å, the remaining 9, 974 micrographs were denoised using the Micrograph Denoiser. 9, 765, 281 particles were picked and extracted with 4x binning from the denoised micrographs using the template picker. The 2D templates were created using EMD-27154. After iterative 2D classifications, 2, 308, 853 pruned particles were subjected to an initial 3D volume (volume 1) reconstruction using 579, 000 particles with *Ab Initio* job as well as four amorphous 3D volumes (volumes 2-5) created by prematurely terminating another *Ab Initio* job. Resulting five 3D volumes were subjected to heterogeneous refinement, and 1, 525, 828 particles belonging to volume 1, the good initial 3D volume, were extracted without binning. Homogeneous refinement was applied to the selected particles to 2.80 Å. Another round of heterogeneous refinement was performed using the 3D volume from the previous homogeneous refinement as 3 repeated input volumes. 1, 158, 228 particles belonging to the resulting classes H0 and H1 were selected and refined to 2.91 Å. These particles were subjected to local and global CTF estimations (41) followed by a non-uniform refinement (42), resulting in a 2.73 Å second reconstruction. A reference-based motion correction was performed on the outputs of the previous non-uniform refinement, and the motion-corrected 1, 157, 747 particles were subjected to another non-uniform refinement, resulting in 2.59 Å reconstruction. 3D flexible refinement (43) was performed on 1, 157, 000 particles, which revealed two dominant molecular motions present in the dataset. To tease out different conformations relevant to the strand displacement activity of Pol γ, local refinement was first performed by creating a focused mask surrounding the catalytic subunit Pol γA and the downstream duplex DNA, then the newly aligned particles were subjected to 3D classification with the same focused mask. Out of 10 classes, 4 classes were selected for further non-uniform refinement, resulting in 2.62 Å, 2.94 Å, 3.23 Å, and 3.05 Å reconstructions for classes C4, C2, C5, and C0, respectively, according to the gold standard FSC (GSFSC) at 0.143.

### Model Building

Cryo-EM maps were post-processed with DeepEMhancer (44) and LocSpiral (45). Pol γ ternary complex structure (PDB: 8D33) was used to build initial models. Manual adjustments were performed in Coot (46) and ISOLDE (47). Atomic models were refined in real-space refinement in Phenix (48, 49) using LocSpiral-processed maps. Refined models were validated with Molprobity and Q-score analysis (49). Structural analysis was performed in Pymol (50) and ChimeraX (51), and structural search was performed with Foldseekn (52). Figures were prepared using ChimeraX.

## Results

### Pol γ strand displacement synthesis under physiological concentrations of Mg^2+^ and Mn^2+^

In many studies evaluating human Pol γ DNA synthesis activity the concentration of divalent metal ion Mg^2+^ is typically from 5 to 20 mM (17, 53, 54)concentrations much higher than the physiological mitochondrial level of 0.45-1.2 mM (55, 56).Considering DNA polymerase (DNAP) activities are sensitive to divalent metal ion concentrations, a report where Pol γ exhibits moderate strand displacement using 4 mM Mg^2+^ and the low Twinkle helicase activity, prompted us to explore Pol γ strand displacement synthesis across a range of 0.02-10 mM Mg^2+^ or Mn^2+^ concentrations using a fork DNA, formed by annealing a 5’-FAM 30-nt primer and an 85-nt flap annealed to a 74-nt template. The fork DNA contained a 40 bp downstream duplex with 4-nt gap and a 45-nt 5’-flap. The same primer and the template were annealed to form p/t for measuring DNA synthesis only (Fig. 1A). Upon addition of dNTPs, Pol γ gap-filling synthesis on the single-stranded template and strand displacement synthesis can be simultaneously monitored by primer extension to 4-nt (P+4) and 44-nt (P+44), respectively.

**Figure 1.**
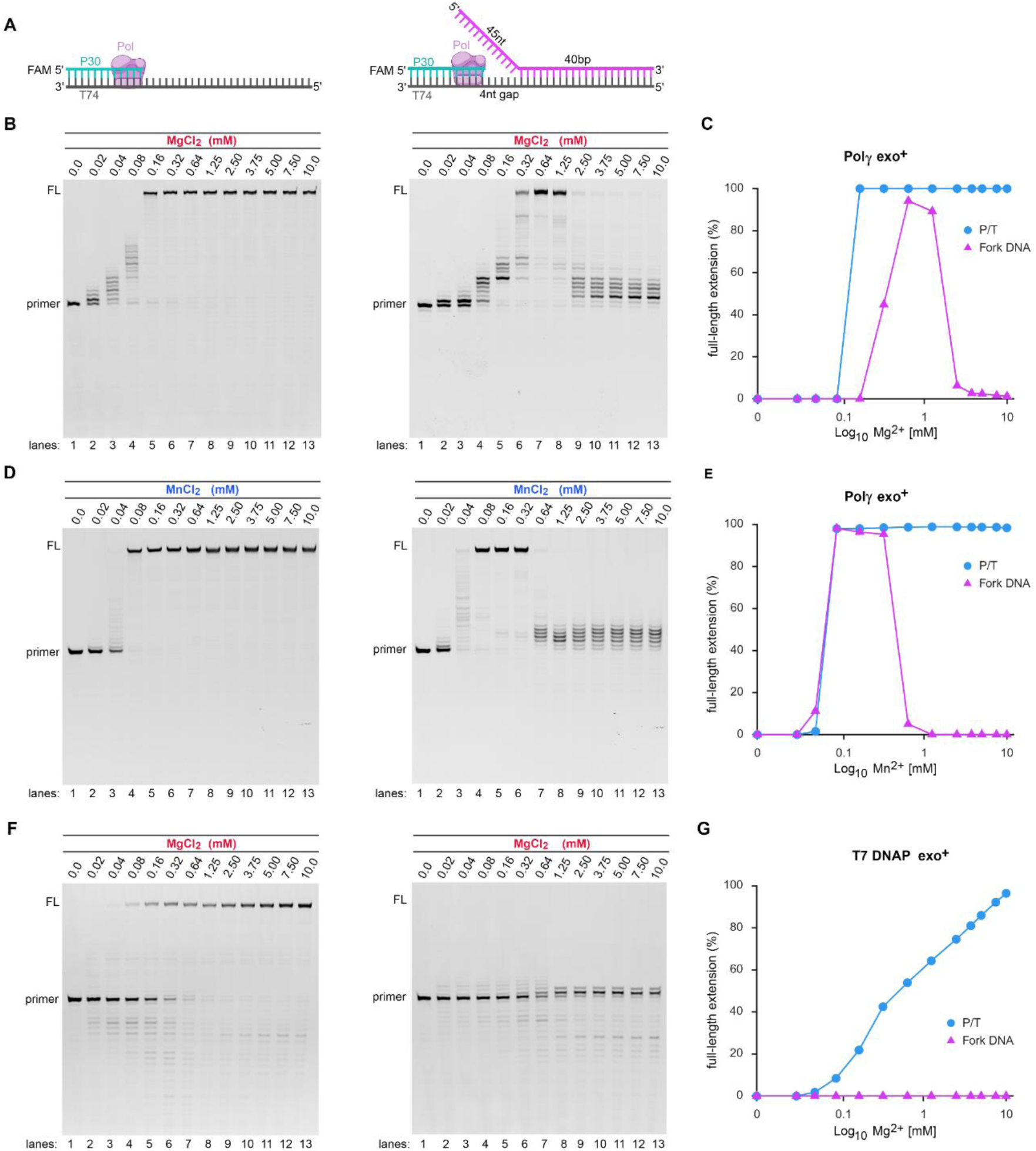
Pol γ metal-dependent DNA strand displacement synthesis activity. (**A**) Schemes of p/t (left) and short fork (right) DNA substrates, (**B**) Pol γ exo+ DNA synthesis on the p/t and fork DNA in the presence of 0 to 10 mM Mg^2+^, (**C**) quantification of percentage of full-length P+44 vs. Mg^2+^ concentration, (**D**) Pol γ Exo+ strand displacement synthesis in the presence of 0 to 10 mM of Mn ^2+^, (**E**) quantification of panel D. (**F**) similar experiments as in panel b, except Pol γ is substituted by T7 DNAP, (**G**) quantification of panel F.

On the single-stranded p/t, Pol γ synthesized products in 10 min with P+44 beginning at 0.02 mM Mg^2+^ and continued up to 10 mM Mg^2+^ equivalently (Fig. 1B, *left panel*). Contrarily, on the fork DNA, Pol γ exhibited strand displacement capability, but only at a narrow [Mg^2+^] range: the product began to form at 0.16 mM Mg^2+^ and reached full length P+44 at 0.32-1.25 mM Mg^2+^, then was absent at Mg^2+^concentrations below 0.16 mM or exceeded 2.5 mM (Fig. 1B, *right panel*). The gap filling synthesis was identical to that on the p/t (Fig. 1B, *right panel*). The results showed that while the *pol* activity is insensitive to Mg^2+^, the DNA unwinding activity only exists at physiological free-Mg^2+^ concentration, and higher Mg^2+^ concentration inhibits Pol γ DNA unwinding (Fig. 1C).

Mn^2+^ is another catalytic metal ion used by DNA polymerases supporting polymerization and exonuclease function. We evaluated whether Mn^2+^ can support Pol γ strand displacement synthesis. Interestingly, on the fork DNA, Pol γ was able to unwind dsDNA generating the P+44 product from 0.08-0.32 mM Mn^2+^ and fell sharply at Mn^2+^ concentrations greater than 0.64 mM (Fig. 1D, *right panel*). Again, Pol γ synthesis on the p/t is insensitive to Mn^2+^ concentration (Fig. 1D, *left panel* and 1E). Thus, Mn^2+^ regulates Pol γ strand displacement at a more restricted concentration than Mg^2+^.

To validate whether Pol γ DNA unwinding is due to the lower metal-ion concentrations weakening downstream dsDNA stability, we first repeated the above assay by using with wild-type T7 DNAP that itself in unable to continue synthesis on a dsDNA (57). If the downstream dsDNA of the fork is weakened at low metal-ion concentration, then T7 DNAP should produce the same metal-dependent strand displacement synthesis. While T7 DNAP formed gap-filled product P+4 at 0.08-10 mM Mg^2+^ on the fork DNA (Fig. 1F, *right*) and P+44 product on the p/t (Fig 1F, *left*), the strand displacement was completely lacking (Fig. 1F, *right panel*), indicating the fork DNA’s downstream 40bp remains duplex (Fig. 1G). We next calculated the melting temperature of the 40bp DNA at various [Mg^2+^] using the Kun’s Oligonucleotide *T*m calculator (58). The *T*m values are 74.4-75.4°C at 0.64-1.25 mM Mg^2+^ (Pol γ SD competent), 76.6-79.1°C at 2.5 mM-10 mM Mg^2+^ and 80.7-82.9°C at 20-50 mM Mg^2+^ (Pol γ SD incompetent) (Table S1). Thus, under our experimental 37°C condition, greater than 99% of downstream DNA exists as a duplex.

### Metal-modulated polymerase tread-milling at the DNA fork junction

Prior studies showed that Pol γ’s reduced strand displacement is due in part to kinetic competition between its polymerase (*pol*) and exonuclease (*exo*) activities at the fork junction-i.e., treadmilling-which prevents forward translocation (24). We tested whether treadmilling is metal dependent using an assay that simultaneously measures *pol* and e*xo* activities. DNA synthesis and excision were quantified on a primed 70-nt circular template (Fig. S1A) after initiating with a dNTP mixture spiked with α-^32^P-dGTP and 0.5–5.0 mM Mg^2+^. Reaction products were separated by thin-layer chromatography to determine the fractions of ^32^P-dGMP incorporated into DNA versus excised. Maximal ^32^P-dGMP incorporation occurred at 1 mM Mg^2+^ (as in Fig. 1) and declined sharply at [Mg^2+^] ≥ 2 mM; conversely, dGMP excision was low at [Mg^2+^] ≤ 1 mM and increased steeply when [Mg^2+^] exceeded 2 mM (Fig. S1B–C). These results indicate that the *exo*/*pol* activity ratio rises with increasing [Mg^2+^], causing the polymerase to idle at the fork and thereby limiting strand displacement.

### Metal sensitivity of Pol γ exo-deficient variants

The above results raise the question of whether Pol γ exo-deficient (exo–) variant, which contains strand displacement activity (24), is regulated by metal-ion concentration. We assayed Pol γ exo–strand displacement synthesis on the same fork DNA (Fig. S2A). Like Pol γ exo+ wild type, the exo– variant exhibited metal-dependent regulation, but over a wider concentration range: maximum full-length products were observed at 0.64–5.0 mM Mg^2+^, declined at 7.5 mM and were diminished at 30 mM Mg^2+^, whereas the *pol* activity on a p/t substrate remains unchanged from 0.32 to 50 mM Mg^2+^ (Fig. S2B-C and S2F). Strikingly, in the presence of Mn^2+^, Pol γ exo– strand displacement synthesis showed metal sensitivity comparable to that of the Pol γ exo+ (Fig. S2D-E and S2G), indicating metal regulation of Pol γ strand displacement is not mediated solely by reduction of exonuclease activity.

### Pol γ proofreading at physiological metal ion concentrations

We next tested whether reduced *exo* activity at low metal ions concentration impacts Pol γ proofreading. The assay was conducted on a primer/template DNA with a T/C mismatched at the 3’-end of the primer (Fig. 2A) (59). If Pol γ proofreads the mismatch and resynthesizes with a correct nucleotide dGTP, a *HindIII* site will be formed; if Pol γ buries the mismatch without proofreading, no *Hind* III site will be formed. The ratio of *Hind* III cleaved vs. noncleaved products was defined as proofreading efficiency. The controls, 40-nt DNA duplexes containing either a mismatch (T/C) or match (G/C) base pair, displayed 0 and 100% proofreading efficiency (Fig. 2B, *lanes* 12-15). In the presence of 0.32 mM, 0.64 mM, 1.25 mM and 5.0 mM Mg^2+^, Pol γ proofreading efficiencies were 88%, 96%, 98% and 100%, respectively (Fig. 2B, *lanes* 4-11 and 2C). The results suggest that the *exo* function is largely intact under conditions that Pol γ is capable of strand displacement synthesis.

**Figure 2.**
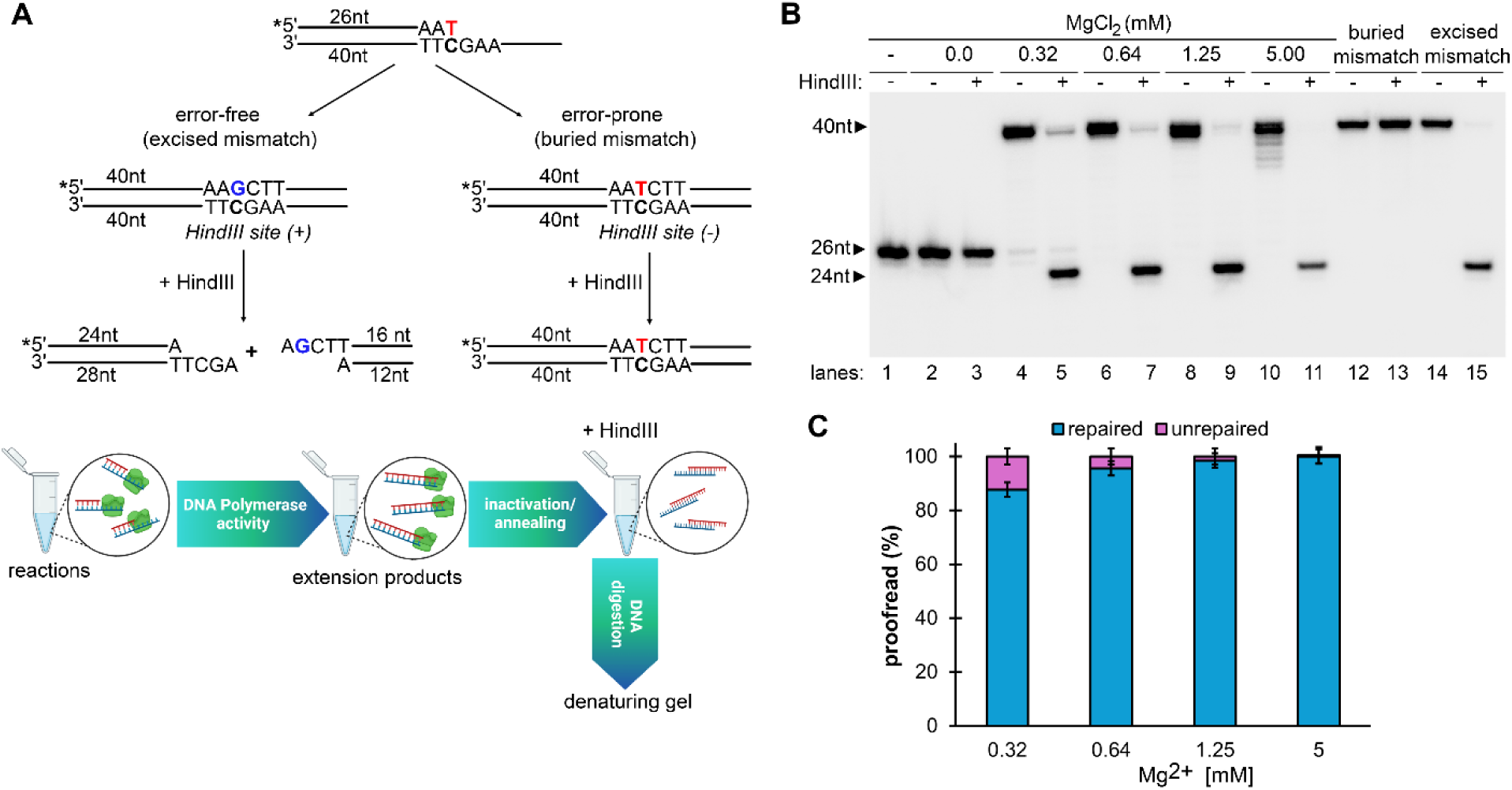
Pol γ mismatch proofreading under different Mg^2+^ concentrations. (**A**) Scheme of experimental design, (**B**) Gel electrophoresis resolved primer extension products with and without Hind III digestion at the indicated Mg^2+^ concentrations. The 26-nt band is the primer, the 24-nt band is a product of Hind III digestion, and (**C**) quantification of percentage of proofread misincorporation at four Mg^2+^ concentrations.

### Pol γ possesses distinct metal binding sites for regulating DNA unwinding

The above results suggest that Pol γ possesses at least two categories of metal binding sites: high (H-) affinity sites that bind to Mg^2+^ at 160 µM-1.25 mM or Mn^2+^ at 80-320 µM, respectively (Fig. 1) and low (L-) affinity sites that bind Mg^2+^ and Mn^2+^ at concentrations greater than 2.5 mM and 0.64 mM, respectively. Metal ions bound in the H-sites support Pol γ simultaneously catalyzing DNA synthesis, proofreading, and dsDNA unwinding, whereas metal ions bound to the L-sites inhibit DNA unwinding.

To sed light on the existence of two distinct-affinity metal binding sites, we tested Pol γ activity under the conditions that the H-and L-sites can be occupied by different metal ions. The assays were carried out at the fixed Mn^2+^ and Mn^2+^ concentration that enabled strand displacement and an increasing concentration of Ca^2+^, commonly used in structural studies as non-reactive cofactor for polymerization and exonuclease reactions (60, 61). If the *pol* activity diminishes, Ca^2+^ has displaced Mn^2+^ in the H-site; if *pol* activity remains and strand displacement diminishes, then Ca^2+^ has occupied the L-sites.

To assess the effects of mixed metal ions on Pol γ activity, replication assays were performed in the presence of a mixture of Ca²⁺ and Mn²⁺ (Ca²⁺/Mn²⁺), a mixture of Ca²⁺ and Mg²⁺ (Ca²⁺/Mg²⁺), as well as Ca²⁺ alone. The Ca²⁺ concentration ranged from 0.02 to 10 mM, while the Mn²⁺ or Mg²⁺ concentration was fixed at levels that support strand displacement synthesis (0.32 mM and 0.64 mM, respectively). To eliminate potential contamination by other metal ions, 1 mM EDTA was added to a 100 mM CaCl₂ stock solution of 99.98% purity.

On the p/t DNA, in the presence of Ca^2+^/Mn^2+^, Pol γ extended the primer to full length in a pattern nearly identical to that obtained with Mn^2+^ alone (compare Fig. 3C with 1D). By contrast, only short products were formed with Ca^2+^ alone (Fig. 3B), indicating that Mn^2+^ remains in the H-sites even in the presence of a 30-fold molar excess of Ca^2+^. Conversely, with Mg^2+^/Ca^2+^, Pol γ activity began to decrease at substoichiometric Ca^2+^/Mg^2+^ molar ratio of 0.5, suggesting that Ca^2+^ occupies the H-sites with a higher affinity than Mg^2+^(Fig. S3A-B).

**Figure 3.**
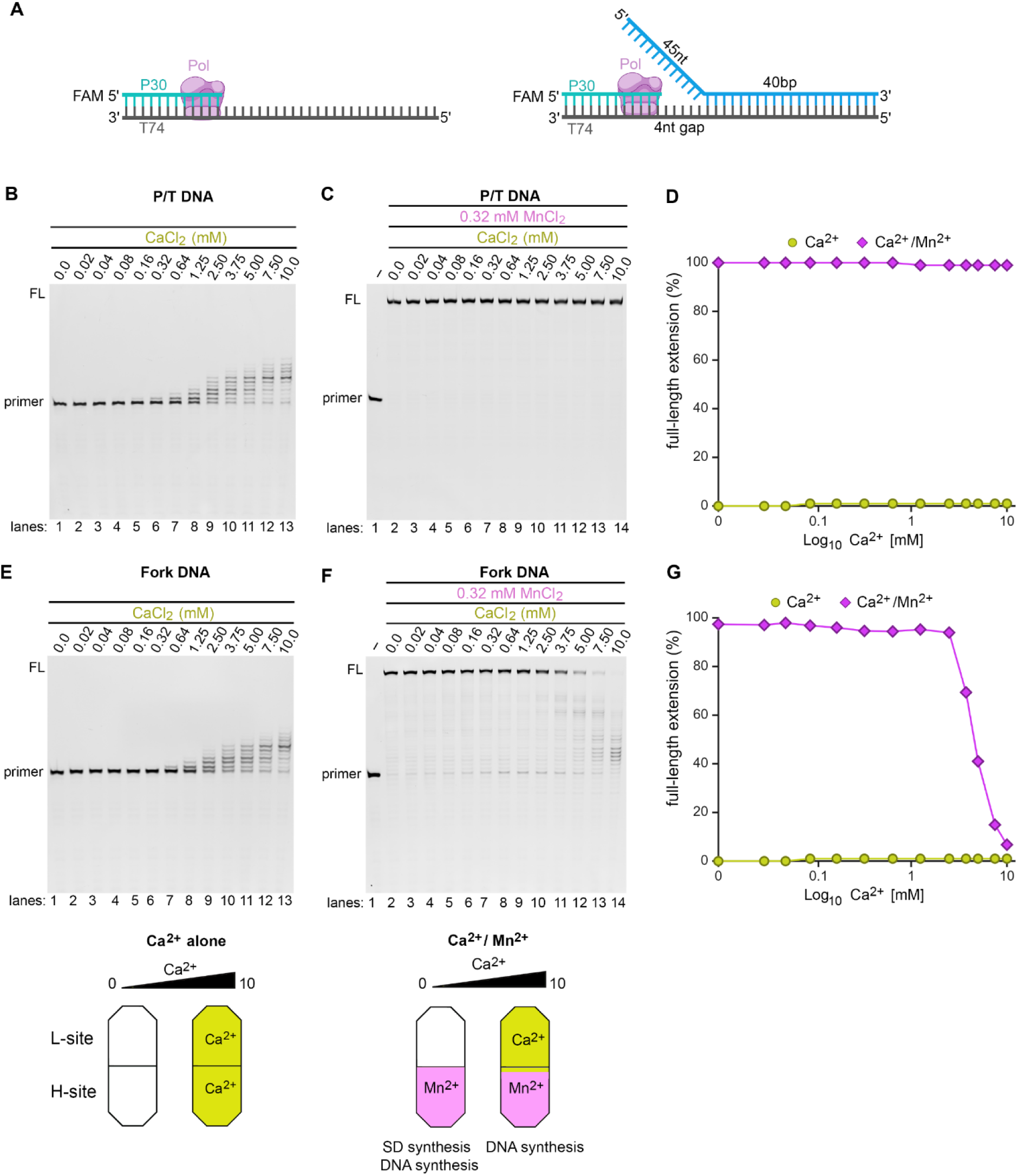
Pol γ strand displacement synthesis in the presence of mixed divalent metal ions. (**A**) p/t and fork DNA substrates used in this assay. (**B**-**D**) Primer extension on a p/t DNA substrate in the presence of 0 to 10 mM Ca^2+^ without (**B**) and with a constant 0.32 mM Mn^2+^ (**C**), (**D**) quantification of b-c panels, (**E-F**) Pol γ SD activity on the fork DNA in the presence of 0 to 10 mM Ca^2+^ without (**E**) and with 0.32 mM Mn^2+^ (**F**), (**G**) quantification of E-F panels.

On the fork DNA, in the presence of Ca^2+^/Mn^2+^, Pol γ strand displacement synthesis was observed at Ca^2+^/Mn^2+^ molar ratio up to 12; and decreased once the ratio exceeded 15 (Fig. 3F and 3G). Since Mn^2+^ remains at the H-site up to molar ratio of 30 (Fig. 3C), the reduced activity is likely due to Ca^2+^ occupying the L-sites, which can also inhibit DNA unwinding. Conversely, in the presence of Ca^2+^/Mg^2+^, strand displacement activity diminished at Ca^2+^/Mg^2+^ molar ratio as low as 0.25 (Fig. S3C)-the same ratio at which Ca^2+^ displaced Mg^2+^ in the H-site (Fig S3D). With Ca^2+^ alone, the products on the fork DNA were nearly identical to that on the p/t (compare Fig. 3E with 3B); Pol γ showed no strand displacement activity with Ca^2+^ alone. These results imply that only Mg^2+^ or Mn^2+^ in the H-site can support strand displacement, whereas occupancy of the L-site by Mg^2+^, Mn^2+^ or Ca^2+^ inhibits DNA unwinding. The experiments suggested affinities of metal ions to the H-sites follow the order of Mn^2+^>Ca^2+^>Mg^2+^, and Mn^2+^>Mg^2+^>Ca^2+^ to the L-sites (Fig. 3D, 3G and S3D).

### Pol γ strand displacement of RNA

The RNA primer that initiates mtDNA replication needs to be removed eventually and resynthesized with DNA. To test Pol γ’s ability to perform the task, we designed a gapped-DNA/RNA construct formed by annealing a 5’-FAM-labeled 30-nt DNA and a 40-nt RNA blocker to the upstream and downstream regions of a 74-nt template, respectively (Fig. 4A). As a control, the same construct was made using a 40-nt DNA blocker. Within 0.32-1.25 mM [Mg^2+^], Pol γ completely displaced RNA and extended all primer to full-length (Fig. 4B); at [Mg^2+^]≥2.5 mM the strand displacement decreases. Similar behavior was observed in the presence of Mn^2+^, where robust RNA displacement synthesis occurred at 0.08-0.32 mM Mn^2+^, then sharply decreased at [Mn^2+^]≥0.64 mM (Fig. 4B-C). We compared the displacement rates of RNA and DNA under the optimal Mg^2+^ concentration for strand displacement, at 0.64 mM. The Pol γ RNA strand displacement rate is comparable to that of the DNA (Fig. 4D-E).

**Figure 4.**
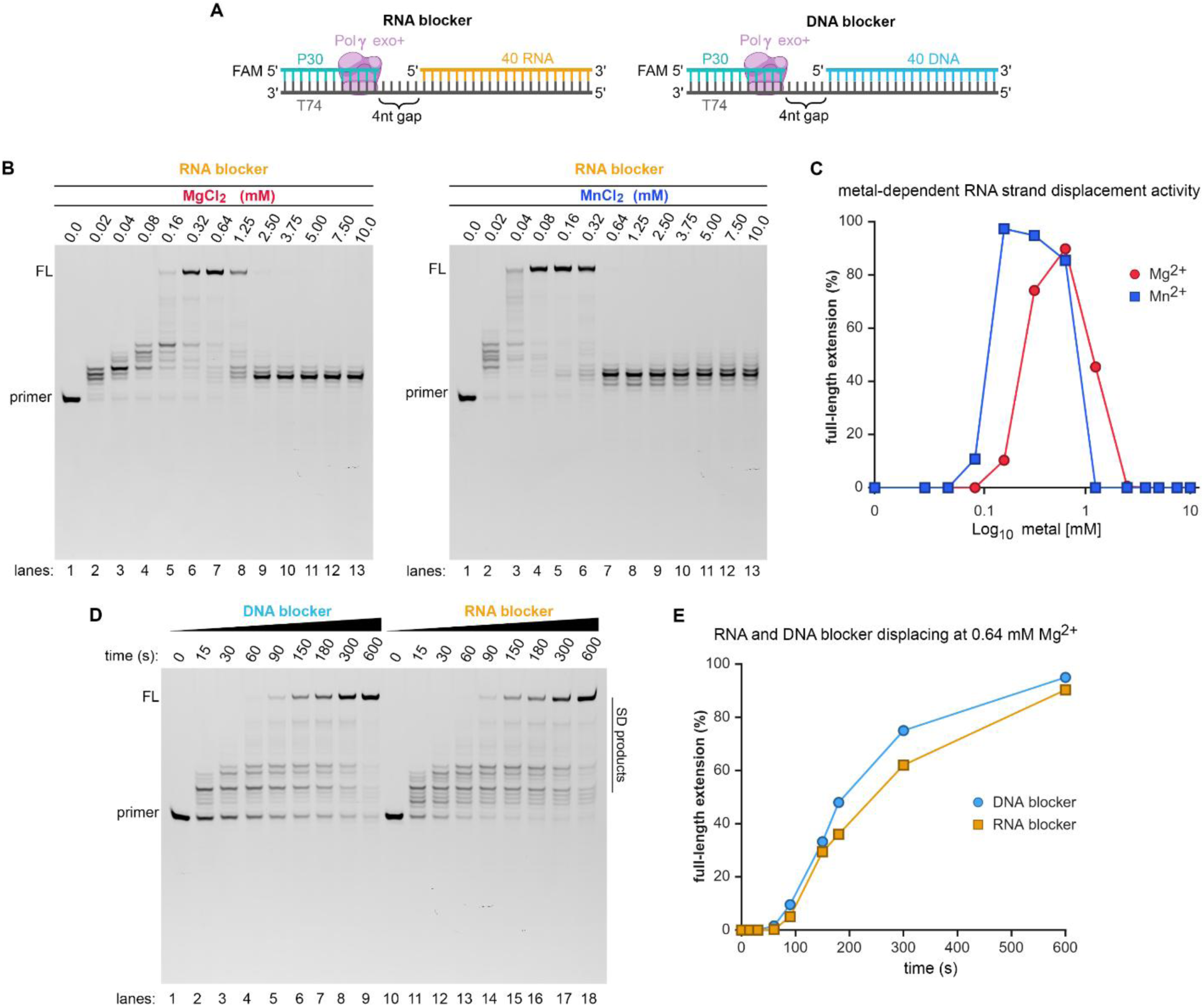
RNA and DNA primer removal by Pol γ exo+. (**A**) scheme of the RNA and DNA blocker substrates used in this assay containing a 4-nt gap and 40-bp downstream region. (**B**) RNA strand displacement assays were performed in a 0 to 10 mM gradient of Mg^2+^ or Mn^2+^ displaying a full-length product (FL) from 0.16-1.25 and 0.04-0.32 mM respectively, (C) Quantification of extended products from panel B. (D-E) Time courses of DNA and RNA displacement syntheses and quantification.

### Localization of metal ion binding sites

In this work, we took advantage of the program MIB2 (62), to predict metal binding sites for Mg^2+^, Mn^2+^ and Ca^2+^ using Pol γ structure as template (PDB 5C51) aiming to find protein region that correlate with the H- and L-sites regulating strand displacement activity. Scores were computed for each amino acid as confidence of its metal binding prediction; a score greater than 1.0 signifies a metal binding site. The prediction accuracy for Mg^2+^, Mn^2+^ and Ca^2+^ was calculated to be 94.6%, 95.0% and 94.1%. Among the Pol γA 1238 amino acids, 144 amino acids are involved in Mg^2+^ binding, 125 in Mn^2+^ binding and 138 in Ca^2+^ binding. For all metal ions, the highest scores are catalytic residues D^890^ and D^1135^ in the *pol* site, D^198^ and E^200^ in the *exo* site. The predictions are consistent with structurally identified metal ion bound to the active sites, lending high confidence to the prediction (Table S2).

Although Pol γ exo– holoenzyme possesses robust strand displacement synthesis capability, the catalytic subunit Pol γA exo– alone does not (16, 17, 24), which suggests Pol γA-Pol γB association is essential for the holoenzyme’s strand displacement. We rationalized that metal ions bound at the subunit interface can probably alter Pol γ holoenzyme strand displacement functions, thus focused on the highly scored Mg^2+^ and Mn^2+^ bound at the Pol γA-Pol γB subunit interface. Three areas are identified: 1) the Pol γA AID subdomain that harbors two predicted binding sites for both Mg^2+^ and Mn^2+^, Glu^538^ and Asp^542^ (Fig. 5B). Glu^538^ forms electrostatic interaction with Pol γB distal monomer R^257^; 2) the Pol γA Thumb subdomain that contains Mg^2+^ potential binding sites at D^465^ and D^469^ that interact with Pol γB proximal monomer at K^373^ and K^365^, respectively (Fig. 5A); 3) and four Mn^2+^ sites at R^443^, E^447^, and E^454^ that interact with PolγB proximal monomer R^264^, R^257^, and D^253^, respectively (Fig. 5A). Binding to positively charged metal ions would weaken the electrostatic interactions at subunit interface, which in turn could reduce holoenzyme strand displacement.

**Figure 5.**
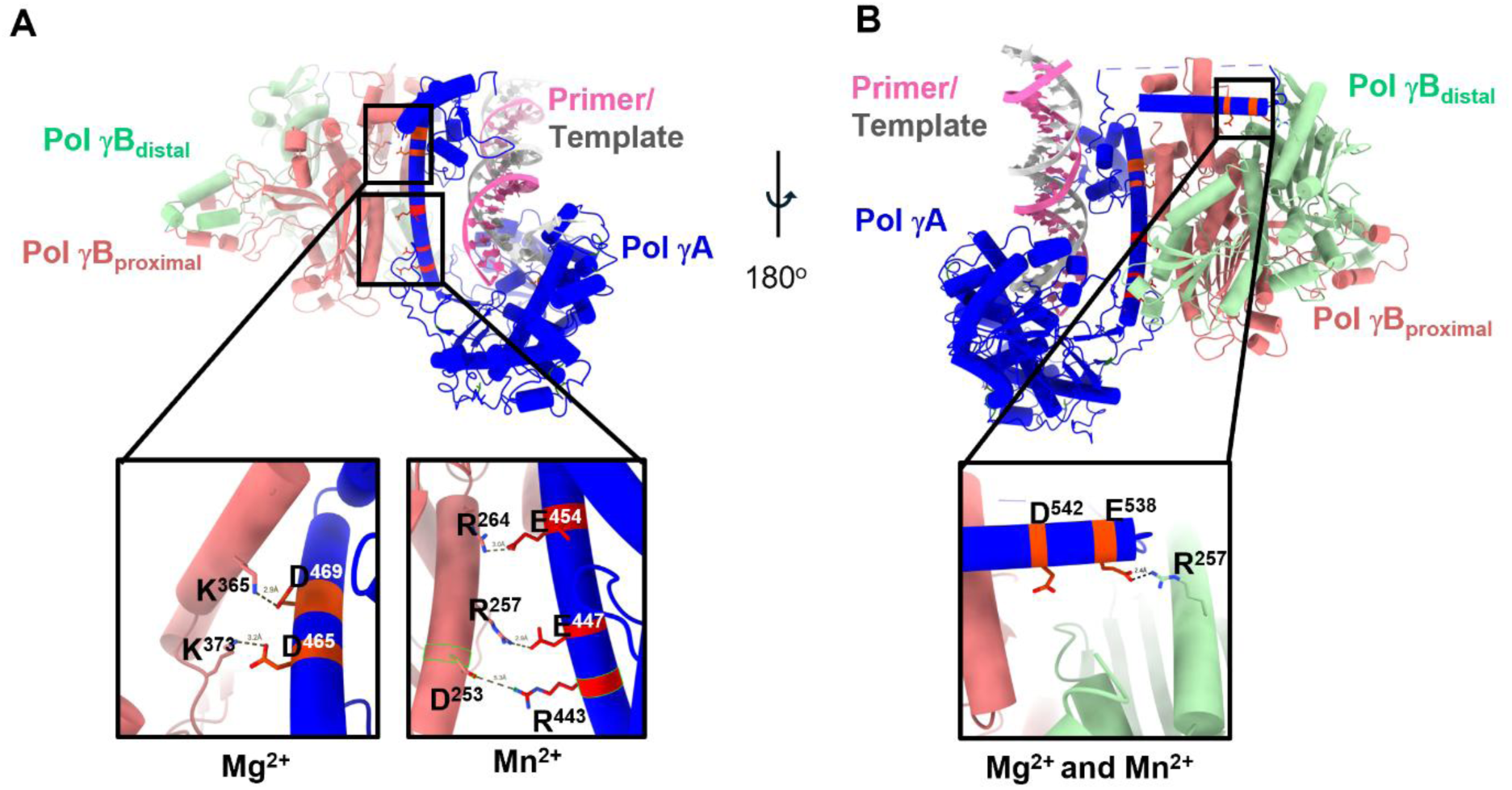
Computationally predicted Mg^2+^ and Mn^2+^ binding sites at subunit interface. Pol γ holoenzyme comprising of Pol γA catalytic subunit (blue) and dimeric Pol γB accessory subunit with proximal (pink) and distal (green) monomers, (**A**) the zoomed-in view of metal-binding sites between the Pol γA Thumb and Pol γB proximal monomer C-terminal domain (CTD), (**B**) metal binding sites between Pol γA AID domain and Pol γB distal monomer CTD.

We carried out dynamic cross-correlation analysis to probe metal ions induced enzyme dynamics. The difference cross-correlation between 20 mM and 1 mM Mg^2+^ showed that metal-concentration induced distinct dynamic coupling and interdomain coordination. At 1 mM Mg^2+^, the Finger domain showed increased anti-correlated motion with the Thumb, Palm or Exo domains (Fig. S4B, purple circles), whereas the Palm and Thumb showed increased correlation with the Exo than the 20 mM Mg^2+^ system (Fig. S4A, green circles).

### Processivity of Pol γ strand displacement activity

Having found that Pol γ has a strong unwinding activity at low Mg^2+^ concentration, the processivity of strand displacement activity was measured on a *midi-rolling circle* template formed by annealing a 5’cy5-71nt primer to a 3.2 kb circular single-stranded DNA annealed (Fig. 6A). The dividing point between primer synthesis on the ss-template and strand displacement synthesis is 3.2 kb. The length of the primer from strand displacement synthesis should represent the polymerase’s processivity, because, unlike on the single-stranded template, the dissociated polymerase is less likely to rebind to the fork junction due to annealing of the template with non-template strands. The assays were conducted in the presence of 0-10 mM Mg^2+^. Like on the short DNA, Polγ synthesis on the 3.2kb ss-template was observed at 0.02-0.08 mM Mg^2+^ where the primer was extended to 1.5 kb in 15 min of reaction. Beginning at 0.16 mM Mg^2+^, strand displacement synthesis was observed, and the maximum primer length reached 6.5 kb at 0.64 mM (Fig. 6B), after which activity decreased. This indicates that the strand displacement synthesis processivity of Pol γ is 3.3-6.5 kb, which is comparable to that on the single-stranded template (63).

**Figure 6.**
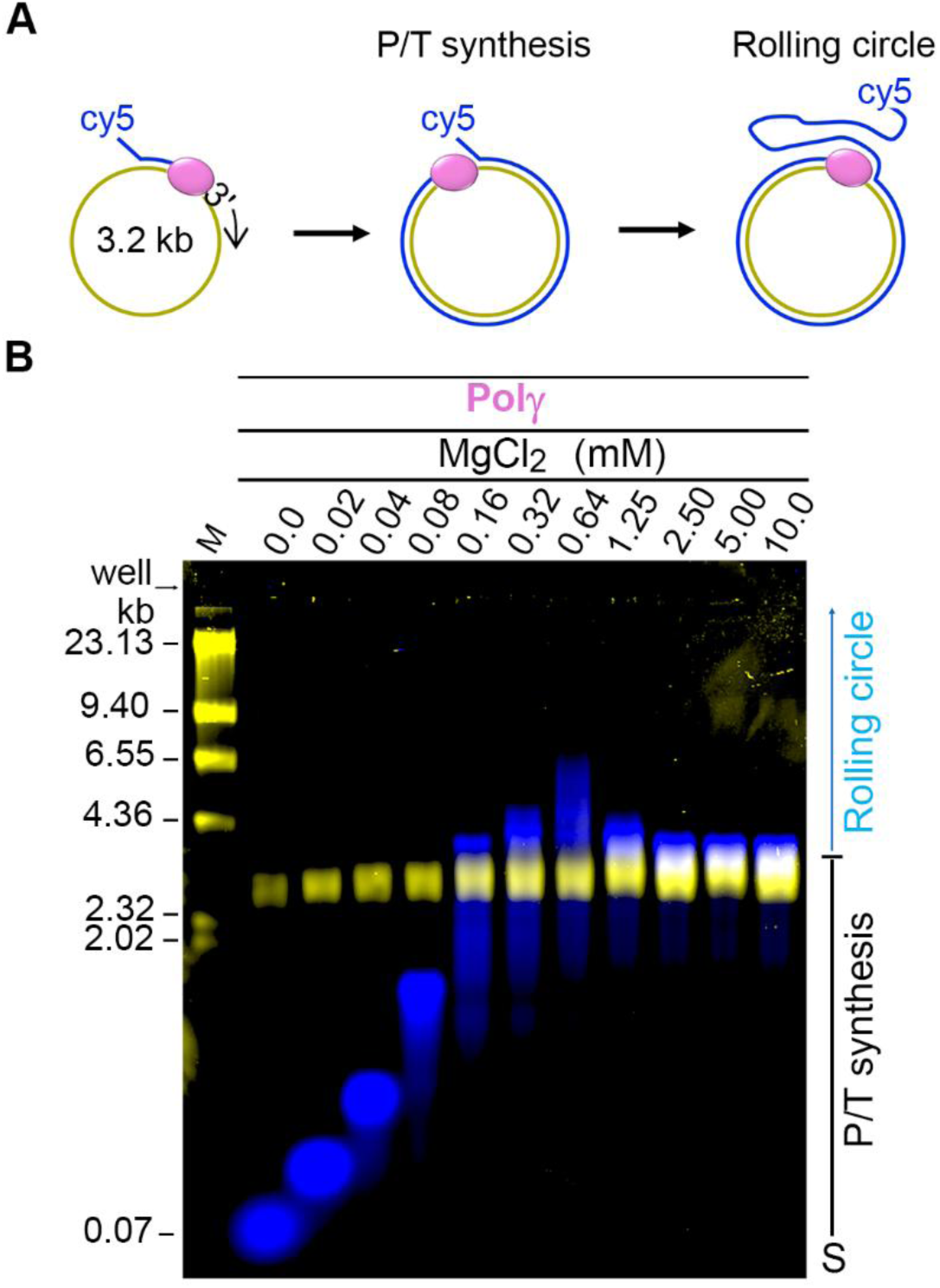
Metal-dependent rolling midi circle synthesis. (**A**) scheme of the midi circle substrate constituted of a 5’-cy5 71-nt primer and a circular ssDNA template, (**B**) products from Pol γ DNA rolling circle synthesis resolved on an alkaline gel (0.8%), scanned using dual-wavelength mode. The circular single-stranded DNA template was visualized with a 488 nm scan for SYBR stain (yellow), and the 5’-cy5-primer and its extension products are visualized with a 635 nm scan (blue).

### Cryo-EM structure of Pol γ-fork DNA complex revealing elements/origin of strand displacement

To elucidate the structural mechanism by which Pol γ unwinds DNA, we determined a cryo-EM structure of the Pol γ ternary complex bound to a 77-nt DNA substrate. The DNA was folded into a dumbbell with a 27-bp upstream and a 7-bp downstream hairpin, separated by a single-nucleotide gap (dT). The primer strand was terminated with 3’-dideoxycytosine (ddC), and the correct incoming nucleotide, dATP, was included (Fig. 7A). Although the enzyme and DNA were of high homogeneity in solution, four cryo-EM structures of Pol γ-DNA complex were dissected to 2.62-3.23 Å resolutions (Fig. 7B-E and Fig. S5). Structural data collection and refinement statistics are shown in Table S3.

**Figure 7.**
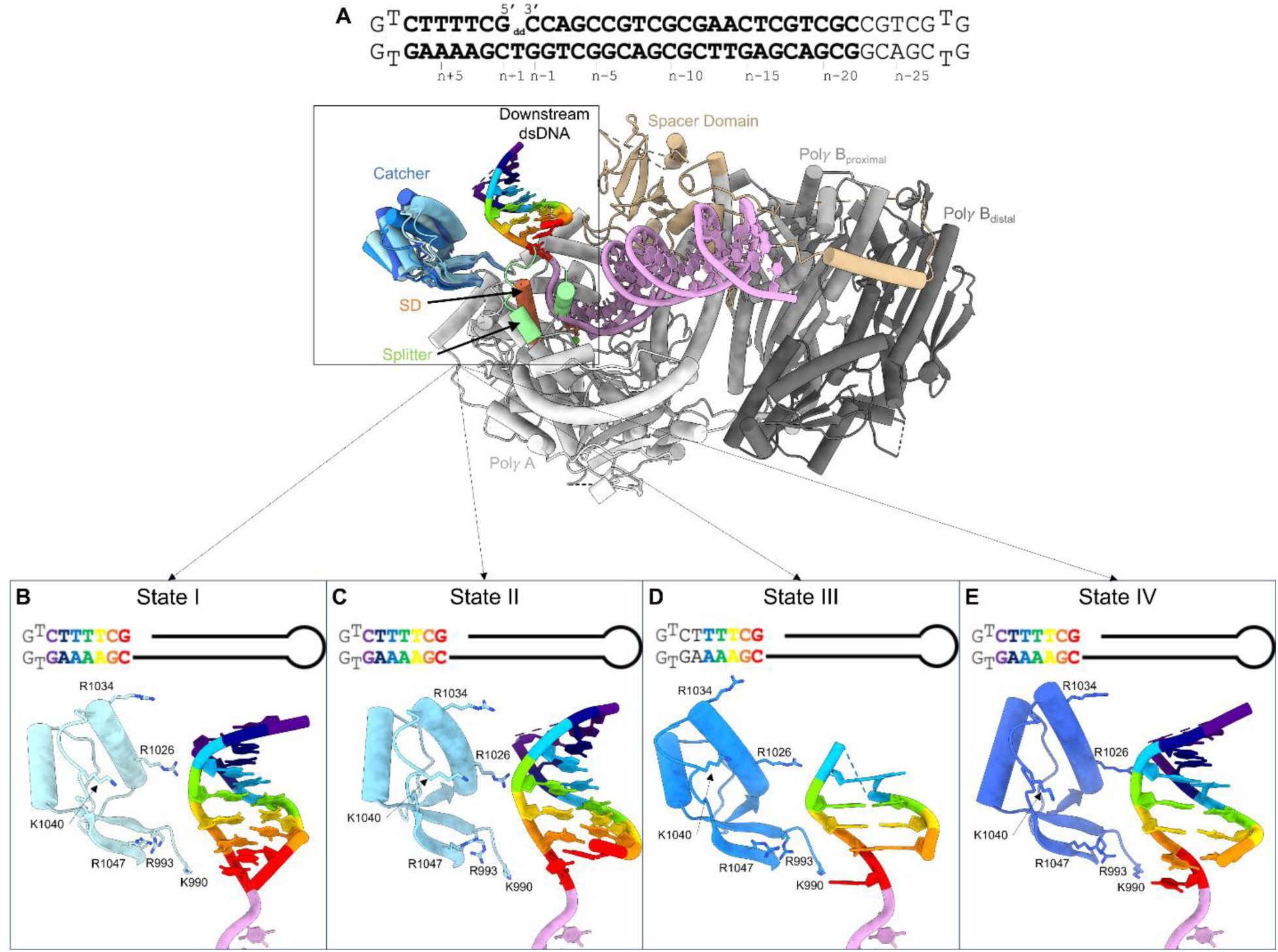
Cryo-EM structures of Pol γ strand displacement complex. (**A**) Dumbbell DNA substrate used in cryo-EM studies. Pol γ-dumbbell DNA complexes superimposed by aligning the Pol site, showing the different conformations of the Catcher domain (different shades of blue). The Catcher, the Splitter and strand displacement (SD) domains are located near the downstream DNA (rainbow color). (**B-E**) close-up views of conformations of the Catcher domain and DNA in different States. (**B**). dsDNA in full-duplex form (State 1), (**C**) the n+1 base pair (bp) buckled (State II), (**D**) the n+1 bp unwound (State III), (**E**) the n+1 bp unwound and the n+2 bp buckled (State IV).

All Pol γ-DNA conformers assumed the replication mode, where the 3’-end of the dumbbell and dATP were in the *pol* site (Fig. S4). While the upstream dsDNA and Pol γ were nearly superimposable, the Catcher domain (Residues 994-1049) and downstream dsDNA exhibited structural diversity (Fig. 7B-E). The downstream 7-bp dsDNA was captured in various unwinding states: a fully base-paired duplex (State I), the buckled *n+1* base pair (State II), the unwound n+1 bp (State III), and the unwound n+1 and buckled *n+2* base pair (IV) (Fig. 8 A-D). Despite different degrees of unwinding, the positions of the template Tn+1 and Tn+2 backbone are invariant, adopting identical positions as those in the ss-template complex (29, 64). The backbone of Tn+1 and Tn+2 rests on a loop (G^956^A^957^G^958^) connecting O- and O1-helices (Fig. S6), stabilizing the dsDNA without specific interaction. The downstream dsDNA in State I-IV exhibited a rolling motion around Tn+1 and Tn+2 of about 20°. As the primer contains a 3’-ddC, no catalysis would occur; the structural ensemble, therefore, represents the dynamics of the downstream duplex at the ground state. The structures showed that the n+1 and n+2 base pairs of downstream dsDNA are less stable and can unwind without any external force. Our cryo-EM results are consistent with observations from NMR studies where 2-3 bp could be spontaneously unwound (65).

**Figure 8.**
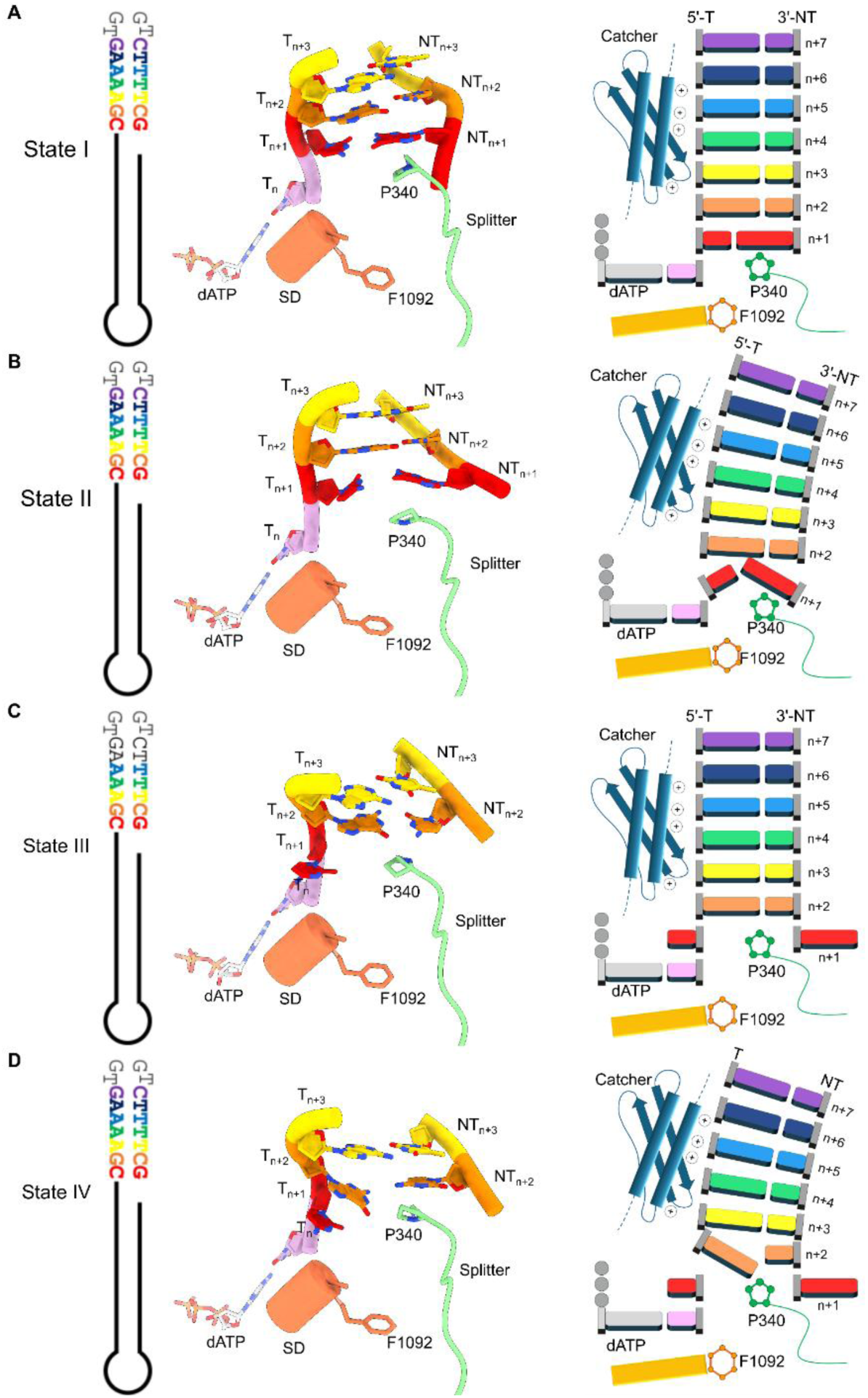
Unwinding mechanism of the downstream dsDNA in Pol γ. (**A-D**) dumbbell DNA substrate with visible downstream duplex shown in corresponding color as the cryo-EM structure (*left*). Pol γ adopts a pre-translocation conformation, where a dATP forms base pair with the template T_n_ in the Pol site (*right*). As no catalysis was permitted, no DNA translocation occurred. Although near the downstream, the SD helix and Splitter loop do not change conformation with the change of dsDNA unwinding state, whereas the Catcher domain undergoes compensatory changes with the dsDNA. In states II and IV, n+1 and n+2 base pairs are buckled against Pro^340^. (*right*) cartoon representation of downstream duplex unwinding along with the correlated movement of the catcher and the downstream duplex.

Three protein elements near the downstream dsDNA fork could potentially participate in DNA unwinding (Fig. 8): The helix-turn-helix Catcher domain (Residues 994-1049) (Fig. 7B-E), which is termed following the nomenclature of yeast mitochondrial DNA polymerase, Mip1 (66), the Splitter helix and the connecting loop (Residues 301-345) that could be involved in proofreading (64), and a Strand Displacement helix (Residues 1087-1100) in the fingers domain (Fig. 7A) (Fig. 7B and 8).

The Catcher domain displayed a polarized charge distribution, the helix facing the downstream DNA is positively charged (Fig. S7C); residues K^1027^, R^1026^, R^1030^, R^1034^, K^1035^, R^1040^ interacting primarily with the template strand of the downstream dsDNA (Fig. 7C-F), and the helix on the opposing side is negatively charged, containing D^999^, E^1000^, E^1002^, and E^1007^ (Fig. S7C). The Catcher domain is less ordered in the Pol γ complexed with a single-stranded template and only emerged in the structure containing dsDNA template. In the dumbbell complexes, the Catcher undergoes compensatory conformational changes with the downstream DNA (Fig. 8, Fig. S7A-B), suggesting their mutual stabilization. Specific electrostatic interactions were seen between R^1049^ and template nucleotide Tn+2 and Tn+3, R^1029^ with template Tn+2 and non-template NTn+3 (Fig. 7B-E). The structure of the Catcher domain is homologous with other DNA/RNA binding proteins, such as *S. cerevisiae* translation machinery-associated protein 64 (RMSD=2.13Å), *B. subtilis* RacA DNA binding protein (RMSD=2.41Å), and *H. sapiens* mdm2 protein (RMSD= 2.49Å) (Fig. S8).

The Splitter is conserved among mitochondrial DNAPs, but the length of connecting loop in human Pol γ is longer than others. Deletion of the W347-L356 in conjunction with A467T was found in patients with Alpers syndrome. Out of 24 children who had the deletion, 21 had Alpers syndrome, comprising hepatoencephalopathy and mtDNA depletion (67).

The tip of the Fingers helix contains a bulky phenylalanine, F^1092^ (Fig. 8). While the helix is slightly off the axis of the downstream DNA, its structural homology to Mip1 and T7 RNAP, together with the flexible downstream duplex location, suggests the helix could still be involved in dsDNA strand separation. The corresponding residue F^848^ in Mip1 was confirmed for the strand displacement activity by mutational studies (66).

Although the above human Pol γ protein elements are potentially involved in strand displacement, we believe that they play passive roles in Pol γ unwinding activity. In Pol γ, the connecting loop of the Splitter helix is located near the n+1 base pair of the downstream duplex. Though partially disordered, the loop’s location implies it could be involved in strand separation. Motifs Pro^340^-Ala^341^ and Asp^756^-Asn^761^ restricted the entrance to the *pol* site to the dimension suitable only for a single-stranded template, thus facilitating strand unwinding while the energy source arises from dNTP binding and incorporation. The presence of the elements might be necessary but insufficient for Pol γ strand displacement synthesis, as the activity is more intricately regulated in the human enzyme (See discussion below).

## Discussion

### Dual roles of Metal ions in Pol γ polymerase and strand displacement activity

Strand displacement synthesis of DNA polymerase (DNAP) refers to concurrent DNA synthesis and dsDNA unwinding, i.e., DNA polymerase and strand displacement activities. The activity is crucial for displacing the RNA primers from Okazaki DNA fragments during lagging strand synthesis, and synthesis on a double-stranded genome without helicase (68, 69). The strand displacement synthesis appears to be idiosyncratic among high-fidelity DNAPs that possess polymerization (*pol*) and exonuclease (*exo*) proofreading activities. Such activity is observed in DNAPs belonging to different structured polymerases, and with varied processivity: Pol A family members DNA Pol I and <u>p</u>lant <u>o</u>rganellar DNA <u>p</u>olymerases (POPs) displayed processivity less than 100 nt (70), whereas the Pol B family bacteriophage phi29 displayed processivity greater than 25 kb (71). Yeast mitochondria Mip1 shows intermediate processivity approximately 4-5 kb (72). Other DNAPs show conditional strand displacement activity. In contrast to their wild-type counterparts, when the *exo* activity is silenced, human Pol δ, Pol ε, Pol γ and bacteriophage T7 DNAP were found to gain strand displacement activity (24, 57, 69, 73–75). Thus, these DNAPs possess intrinsic strand-displacement capability, and the balance between *pol* and *exo* activities regulates strand displacement activity.

Previous studies in E. coli DNAP I have shown the existence of additional metal binding sites different from the canonical *pol* and e*xo* catalytic sites associated with polymerization regulation (76). In the study they found tight sites corresponding to the exonuclease and polymerization active sites and about 20 weak sites which are located surrounding these domains but out of the catalytic place. Here, we identified another regulatorof Pol γ strand displacement synthesis, divalent metal ions. We propose that Pol γ possesses two classes of metal binding sites with distinct affinities that independently control DNA synthesis and unwinding. High-affinity (H) sites within the *pol* and *exo* active sites support DNA synthesis and enable DNA unwinding, whereas the low-affinity (L) sites, outside of active sites, suppress DNA unwinding without affecting DNA synthesis (Fig. 9). Only catalytically metal, Mg^2+^ and Mn^2+^, in the H-sites can support strand displacement synthesis, while occupancy of the L-sites by Mg^2+^, Mn^2+^ or catalytically incompetent Ca^2+^ inhibits DNA unwinding (Fig. 9). At physiological concentrations of metal ions, the H-sites are saturated, allowing Pol γ to carry out DNA synthesis and DNA unwinding simultaneously. We further show that Pol γ unwinds RNA/DNA hybrid with equal efficiency as DNA/DNA duplex, suggesting a role in RNA primers removal.

**Figure 9.**
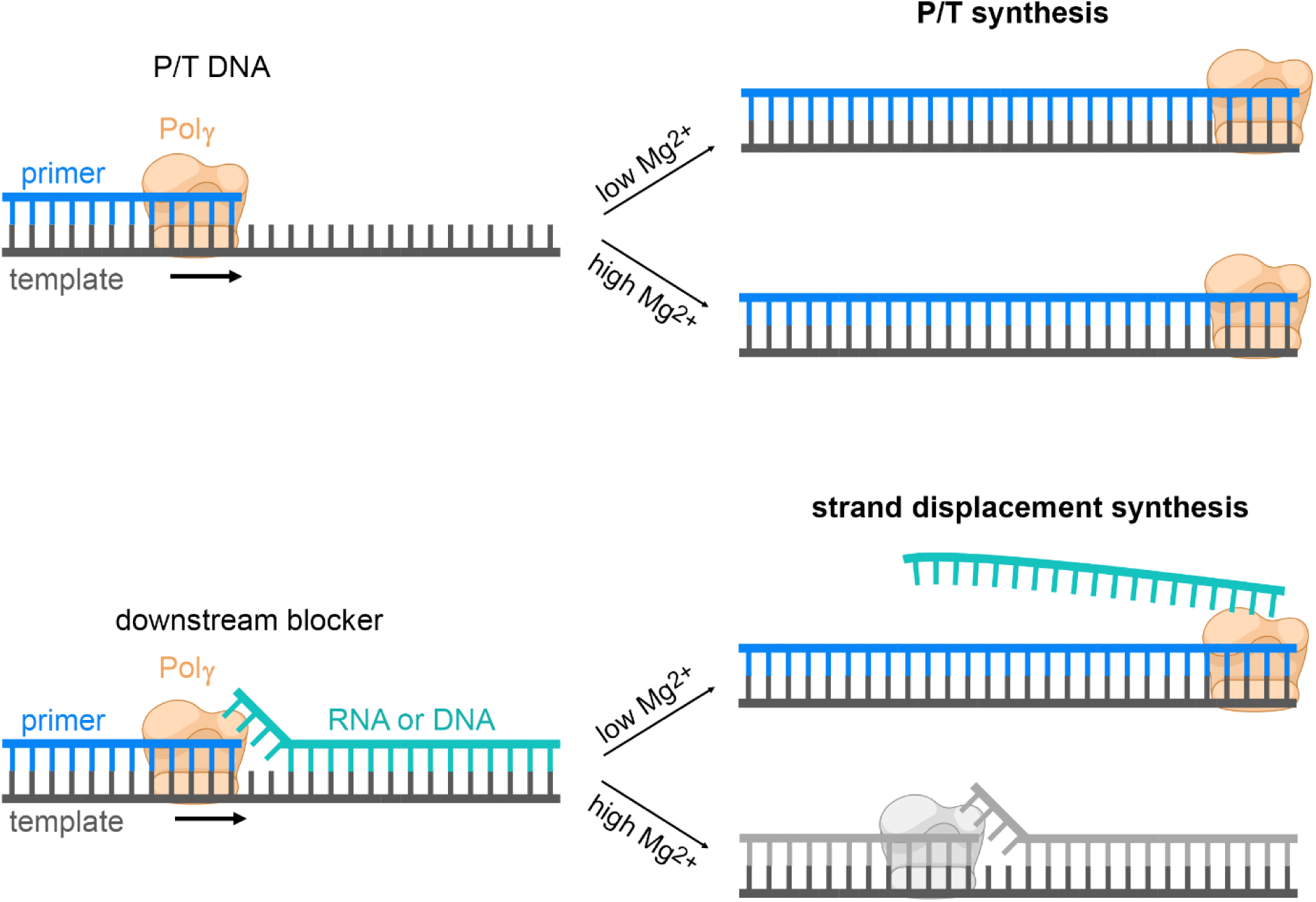
Mechanism of the metal-regulated Pol γ strand displacement synthesis. Extension of p/t DNA is not affected by high Mg2+ concentration. Low Mg2+ concentration allows RNA or DNA strand displacement while high concentrations become inhibitory.

The metal-regulated strand displacement is not strictly via *exo* activity, as the strand displacement of Pol γ exo– variant is also sensitive to divalent metal ion concentrations, although the regulation occurs at higher Mg^2+^ concentration than that for the Pol γ exo+. While DNA unwinding of Pol γ exo– showed comparable sensitivity to Mn^2+^ as the Pol γ exo+. We showed that under the condition that Pol γ can carry out strand displacement, its *exo* function is intact. Thus, regulation of strand displacement by metal ions does not share the pathway of the *exo* activity.

The mechanism for metal regulation of Pol γ strand displacement synthesis stems from the structure and composition of the holoenzyme. Pol γ holoenzyme consists of a catalytic subunit Pol γA and an accessory subunit Pol γB. While Pol γ exo– variant displays strong strand displacement capability, Pol γA exo– has been shown to unwind in a limited manner, indicating interaction between Pol γA and Pol γB is essential for strand displacement. Ourcomputational analysis showed high-confidence metal binding sites in two Pol γA-Pol γB interfaces, which could disrupt electrostatic interaction of the subunits and weakens the holoenzyme (Fig. 9, bottom).

Molecular dynamics simulations indicate that Pol γ exhibits significantly greater structural rigidification at high Mg²⁺ concentrations (20 mM) than the lower concentration (1 mM), with reduced root-mean-square-fluctuations in the Palm, Thumb and Fingers domains, suggesting that metal ions in the L-sites increase the rigidity Pol γ that is unfavorable for strand displacement activity but does not impair the polymerase activity.

### Structural and molecular model of DNA polymerase strand-displacement synthesis

DNAPs that possess strand displacement synthesis activity are found in both the Pol A and Pol B families (77). Upon examination, we observed that their structural features near the downstream DNA entry site are diverse. Among Pol A family members (78)—including Pol γ, Mip1, DNA Pol I, and T7 DNAP—the common feature is a “C-shaped” *pol*-site entrance that cradles the template strand while displacing the non-template strand (Fig. 10, top). In the Pol B family members (79)—including Pol δ, Pol ε, and RB69— a similar ‘C-shape’ structure was also observed. Phi29 DNAP, which exhibits the strongest strand displacement activity, forms a closed “O-shaped” *pol*-site entrance that fully encircles the template strand and completely excludes the non-template strand (71, 80) (Fig. 10, bottom).

**Figure 10.**
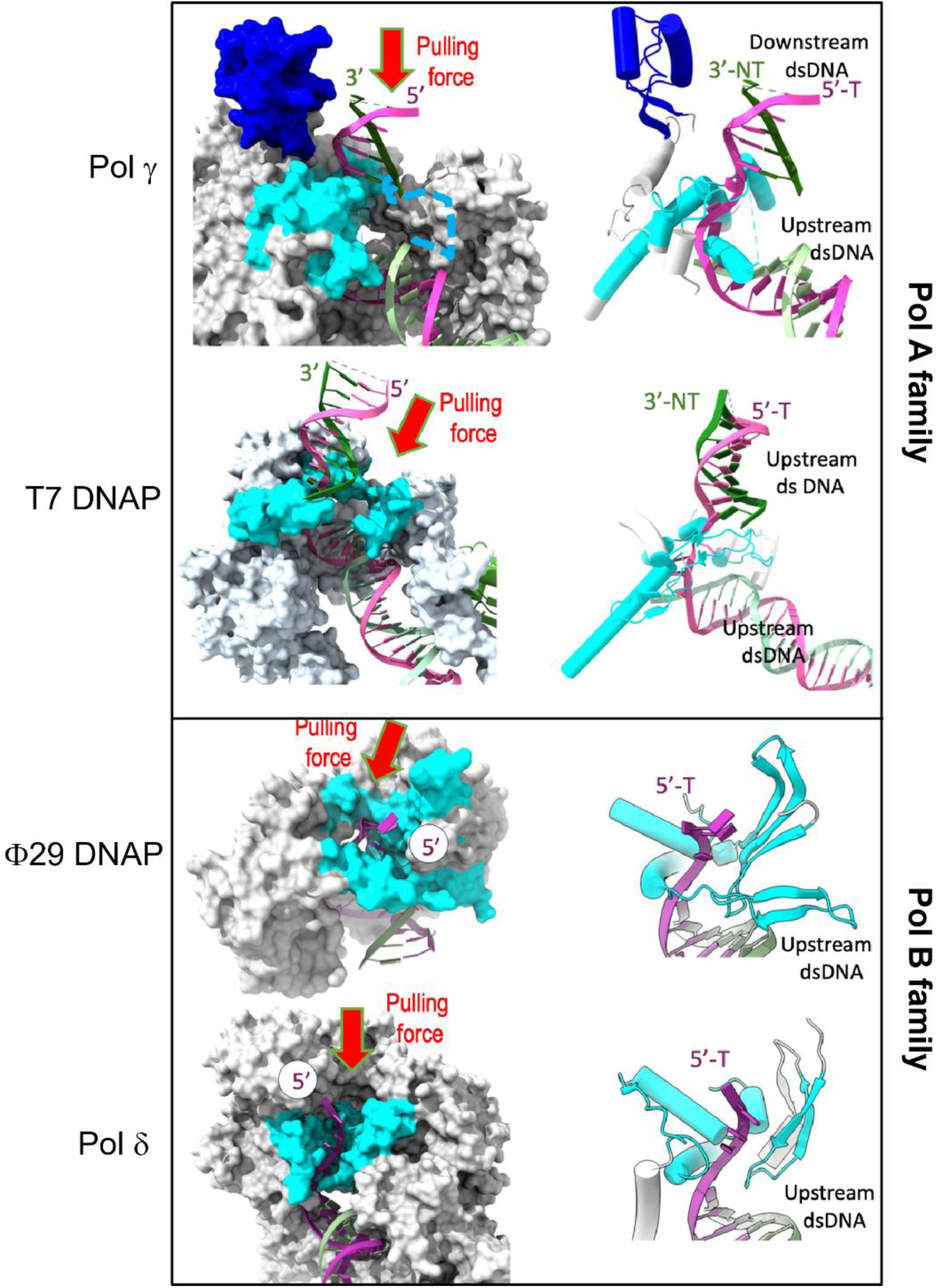
Diverse structural features of DNAPs with strand displacement capability at the Pol site entrance. Pol A family members, Pol γ and T7 DNAP, showed a ‘C-shaped’ Pol site entrance that facilitates displacement of the non-template (NT) strand, while the corresponding protein elements are not conserved. The dashed line in Pol γ represents thedisordered portion of the Splitter domain, whose AlphaFold predicted structure showed high diversity (Fig. S7). Pol B family members, phi29 DNAP and Pol δ. The Pol site entrance in phi29 DNAP is completely enclosed, forming an ‘O-shape, while in Pol δ is ‘C-shape’, and the structural components of the entrance are not strictly conserved.

However, the structural composition of the *pol*-site entrance is not conserved either within or across these families. In Pol A family, T7 DNAP, which contains the SD helix, has a unique positively charged β-turn-β hairpin (residues 575-590) as a part of the ‘C-shaped’ entrance. Substitutions of the hairpin’s basic K587, K589, R590, and R591 with charge-neutral Asp in T7 phage resulted in reduced DNA synthesis and a 10-fold decrease in growth rate (81), underscoring the functional importance of this hairpin. The corresponding position of the T7 DNAP hairpin is occupied by a helix in DNA Pol I (82), and by a portion of the Splitter in Pol γ, and is absent in Mip1 (66) (Fig.10). In Pol B family, the *pol* entrance consists of β-sheets and α-helices, which share no structural homology with the Pol A family members. The helices contact the template strand of the downstream dsDNA, whereas the β-sheets would contact the non-template strand. In RB69 DNAP (83) and Pol δ (84), only a single pair of β-sheets is observed. In contrast, the same position in phi29 DNAP is occupied by a triple β-sheet structure, furthermore, Phi29 DNAP contains an additional pair of β-sheets that fully encloses the *pol*-site entrance (Fig.10).

Based on these observations, we developed a model for DNAP strand displacement synthesis. In this model, the energy driving unwinding arises from nucleotide incorporation, while the strand displacement elements surrounding the dsDNA restrict the dimensions of the template strand’s entry into the *pol*-site. Each dNTP incorporation provides ∼5–7 kcal/mol of binding free energy plus ∼7.3 kcal/mol from hydrolysis to dNMP + PPi, yielding >10 kcal/mol of catalytic energy (85–87). This is sufficient to unwind a base pair (2–3 kcal/mol for AT or GC pairs). After nucleotide incorporation, polymerase translocation generates a “Pulling Force” (Fig. 10) on the template strand through the physically constrained *pol*-site entrance, which only allows a single-stranded template to pass. As a result, the non-template strand is displaced.

In conclusion, our study provides a structural and functional basis for human Pol γ strand displacement synthesis and regulation of this activity by metal ions. Divalent metal ions Mg^2+^ and Mn^2+^ play a dual role in Pol γ DNA synthesis catalysis and regulation of dsDNA strand separation. Under physiological concentrations of metal ions, Pol γ has robust strand displacement capability with high processivity.

## Data availability

Cryo-EM density maps and atomic coordinates have been deposited using the Worldwide Protein Data Bank (wwPDB) OneDep System. Pol γ strand displacement ternary-complexes are available under the following accession codes: State I (EMD-72478) (PDB: 9Y4C), State II (EMD-72479) (PDB: 9Y4D), State III (EMD-72480) (PDB: 9Y4E), and State IV (EMD-72481) (PDB: 9Y4F).

## Supplementary Data Statement

Supplementary Data are available once published

## Author contributions Statement

N.B.T and Y.W.Y. conceived the project. N.B.T. performed biochemical experiments, protein purification and data analysis. J.P. performed data collection and structural determination. J.M-G and L.G.B. obtained the midi circle DNA substrate and provided reaction conditions starting points. A.R and A.C performed *in silico* simulation and data interpretation. A.S performed biochemical assay and analysis. S.S.P. and Y.W.Y. supervised the project. N.B.T. and Y.W.Y. wrote the first draft, and all authors contributed to the final manuscript.

## Funding

The work is supported by NIH grants (R01AI134611 and R01GM145925 to Y.W.Y., MIRA GM118086 to S.S.P., and R35GM151951 to G.A.C.). Latin American Fellowship in Biomedical Sciences and supported in part by the Pew Charitable Trusts to N.B.T., the Jeane B. Kempner postdoctoral fellowship to J.P., and an endowment from the Sealy and Smith Foundation to the Sealy Center for Structural Biology and Molecular Biophysics at UTMB.

## Conflict of Interest

The authors declare no competing interests.

## Supplemental Information

**Table S1.**
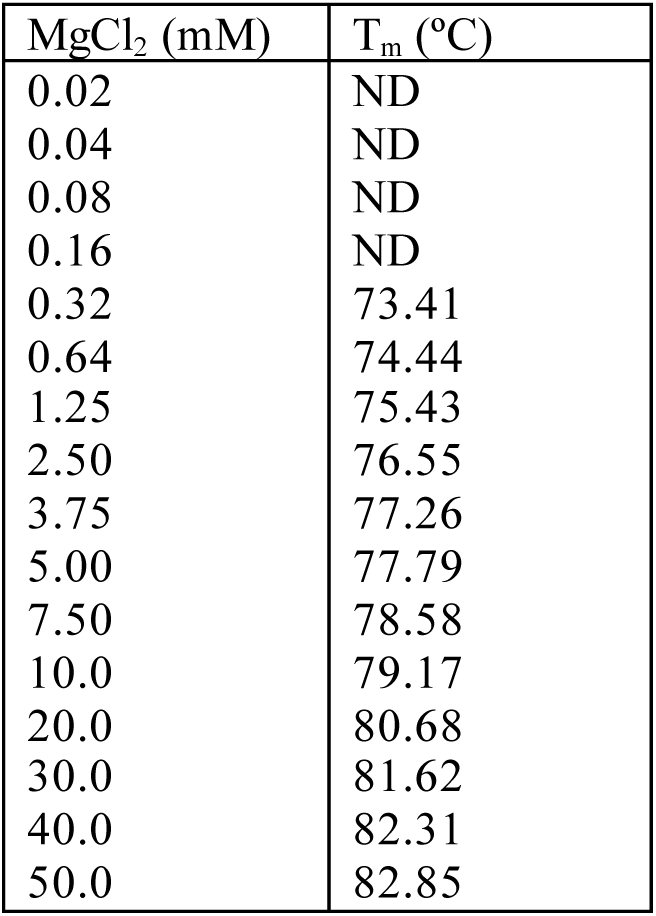
Tm values for dsDNA at different Mg^2+^ concentrations.

The following parameters were used: 140 mM NaCl, 0.2 mM dNTPs, and 100 nM the fork DNA substrate (40-bp duplex region). Data obtained using Kun’s Oligonucleotide Tm calculator (https://arep.med.harvard.edu/kzhang/cgi-bin/myOligoTm.cgi) assuming 99.9% of annealed template.

**Table S2.** Pol γ metal binding sites predicted by MIB2 using structure (PDB 5C51).

| Rank | **Mg<sup>2+</sup>** |  |  | **Mn<sup>2+</sup>** |  |  | **Ca<sup>2+</sup>** |  |  |
| --- | --- | --- | --- | --- | --- | --- | --- | --- | --- |
|  | Res # | Res | Score | Res # | Res | Score | Res # | Res | Score |
| 1 | **890** | **ASP** | **4.39** | 453 | GLU | 5.99 | **1135** | **ASP** | **5.14** |
| 2 | **1135** | **ASP** | **4.39** | 454 | GLU | 5.99 | 1136 | GLU | 5.14 |
| 3 | 1136 | GLU | 4.39 | 457 | ARG | 5.99 | **890** | **ASP** | **4.93** |
| 4 | 891 | VAL | 3.56 | **890** | **ASP** | **4.85** | 891 | VAL | 4.93 |
| 5 | 892 | ASP | 3.24 | **1135** | **ASP** | **4.85** | 893 | SER | 3.88 |
| 6 | 1186 | ASP | 3.19 | 1136 | GLU | 4.48 | 892 | ASP | 3.54 |
| 7 | 1187 | ARG | 3.19 | 1186 | ASP | 3.68 | **198** | **ASP** | **3.20** |
| 8 | 134 | ASN | 3.11 | 1192 | GLU | 3.68 | 199 | VAL | 3.20 |
| 9 | 136 | ASP | 3.11 | 1196 | ASP | 3.68 | **200** | **GLU** | **3.20** |
| 10 | 1068 | ASP | 2.97 | 892 | ASP | 3.40 | 536 | GLU | 2.92 |
| 11 | 1084 | GLU | 2.97 | 358 | GLU | 3.37 | 537 | GLU | 2.92 |
| 12 | 1085 | PRO | 2.97 | 361 | ARG | 3.37 | 489 | GLU | 2.79 |
| 13 | 122 | ASP | 2.94 | 930 | ASP | 3.34 | 491 | ASP | 2.79 |
| 14 | 123 | VAL | 2.94 | 932 | HIS | 3.34 | 928 | GLY | 2.68 |
| 15 | 933 | SER | 2.59 | 773 | LYS | 3.22 | 930 | ASP | 2.68 |
| 16 | 934 | LYS | 2.59 | 776 | ASP | 3.22 | 932 | HIS | 2.68 |
| 17 | 936 | ALA | 2.59 | 136 | ASP | 2.98 | 136 | ASP | 2.68 |
| 18 | 346 | ASP | 2.40 | 140 | ARG | 2.98 | 137 | GLN | 2.68 |
| 19 | 349 | ASP | 2.40 | 1134 | HIS | 2.89 | 832 | ASP | 2.56 |
| 20 | 350 | ILE | 2.40 | 241 | PRO | 2.88 | 833 | GLU | 2.56 |
| 21 | 1081 | ARG | 2.40 | 243 | ASP | 2.88 | 834 | GLU | 2.56 |
| 22 | **198** | **ASP** | 2.38 | 244 | LEU | 2.88 | 465 | ASP | 2.40 |
| 23 | 199 | VAL | 2.38 | 1188 | CYS | 2.69 | 468 | ASN | 2.40 |
| 24 | **200** | **GLU** | 2.38 | 1215 | GLY | 2.69 | 469 | ASP | 2.40 |
| 25 | 399 | ASP | 2.38 | 443 | ARG | 2.67 | 925 | LYS | 2.35 |
| 26 | 951 | TYR | 2.34 | 447 | GLU | 2.67 | 1182 | ALA | 2.35 |
| 27 | 952 | GLY | 2.34 | 538 | GLU | 2.63 | 346 | ASP | 2.26 |
| 28 | 538 | GLU | 2.31 | 542 | ASP | 2.63 | 349 | ASP | 2.26 |
| 29 | 542 | ASP | 2.31 | 925 | LYS | 2.63 | 350 | ILE | 2.26 |
| 30 | 1172 | ASP | 2.26 | 891 | VAL | 2.39 | 895 | GLU | 2.17 |
| 31 | 1173 | LEU | 2.26 | 1103 | SER | 2.20 | 894 | GLN | 2.11 |
| 32 | 1182 | ALA | 2.26 | 1107 | ASP | 2.20 | 1134 | HIS | 2.11 |
| 33 | 293 | ASP | 2.18 | 346 | ASP | 2.15 | 384 | ASP | 2.08 |
| 34 | 294 | THR | 2.18 | 349 | ASP | 2.15 | 387 | GLU | 2.08 |
| 35 | 894 | GLN | 2.16 | 350 | ILE | 2.15 | 388 | ASN | 2.08 |
| 36 | 832 | ASP | 2.13 | 1081 | ARG | 2.15 | 1133 | ILE | 2.06 |
| 37 | 833 | GLU | 2.13 | 1126 | ASP | 2.13 | 145 | LYS | 2.00 |
| 38 | 834 | GLU | 2.13 | 1145 | ASP | 2.13 | 146 | GLN | 2.00 |
| 39 | 1219 | ASP | 2.08 | 1148 | ARG | 2.13 | 147 | SER | 2.00 |
| 40 | 1222 | GLN | 2.08 | **198** | **ASP** | **2.06** | 1068 | ASP | 1.94 |
| 41 | 1223 | ILE | 2.08 | **200** | **GLU** | **2.06** | 1069 | ILE | 1.94 |
| 42 | 1225 | GLU | 2.07 | 399 | ASP | 2.06 | 72 | ASN | 1.91 |
| 43 | 1226 | LEU | 2.07 | 928 | GLY | 1.95 | 75 | ASP | 1.91 |
| 44 | 1107 | ASP | 2.06 | 404 | HIS | 1.92 | 1191 | LYS | 1.82 |
| 45 | 1110 | HIS | 2.06 | 405 | GLU | 1.92 | 776 | ASP | 1.81 |
| 46 | 830 | ASP | 2.03 | 1191 | LYS | 1.81 | 777 | GLY | 1.81 |
| 47 | 465 | ASP | 2.02 | 1181 | SER | 1.80 | 903 | GLY | 1.81 |
| 48 | 468 | ASN | 2.02 | 615 | SER | 1.79 | 904 | ASP | 1.81 |
| 49 | 469 | ASP | 2.02 | 741 | ASP | 1.79 | 1219 | ASP | 1.78 |
| 50 | 274 | ASP | 2.01 | 743 | ASP | 1.79 | 1220 | ILE | 1.78 |
| 51 | 466 | LEU | 1.95 | 489 | GLU | **1.78** | 579 | ARG | 1.78 |
| 52 | 900 | ALA | 1.92 | 491 | ASP | 1.78 | 581 | ASP | 1.78 |
| 53 | 904 | ASP | 1.92 | 943 | ARG | 1.76 | 582 | ASP | 1.78 |
| 54 | 1067 | SER | 1.92 | 1193 | VAL | 1.71 | 743 | ASP | 1.67 |
| 55 | 895 | GLU | 1.91 | 828 | HIS | 1.68 | 744 | ILE | 1.67 |
| 56 | 902 | LEU | 1.90 | 1143 | GLU | 1.68 | 746 | GLY | 1.67 |
| 57 | 903 | GLY | 1.90 | 1056 | GLU | 1.66 | 749 | PHE | 1.67 |
| 58 | 429 | GLU | 1.87 | 1059 | ASN | 1.66 | 398 | GLN | 1.66 |
| 59 | 1117 | LYS | 1.87 | 933 | SER | 1.64 | 399 | ASP | 1.66 |
| 60 | 1191 | LYS | 1.80 | 226 | GLN | 1.58 | 608 | ASP | 1.64 |
| 61 | 893 | SER | 1.79 | 230 | GLU | 1.58 | 1082 | ALA | 1.60 |
| 62 | 1069 | ILE | 1.75 | 756 | ASP | 1.55 | 372 | GLU | 1.57 |
| 63 | 407 | PHE | 1.70 | 761 | ASN | 1.55 | 373 | PRO | 1.57 |
| 64 | 408 | GLN | 1.70 | 1190 | ARG | 1.52 | 375 | GLU | 1.57 |
| 65 | 1165 | ALA | 1.70 | 1138 | ARG | 1.52 | 391 | ASP | 1.57 |
| 66 | 1166 | TYR | 1.70 | 777 | GLY | 1.50 | 269 | HIS | 1.57 |
| 67 | 1080 | SER | 1.60 | 894 | GLN | 1.48 | 274 | ASP | 1.57 |
| 68 | 1082 | ALA | 1.60 | 370 | GLU | 1.46 | 122 | ASP | 1.54 |
| 69 | 226 | GLN | 1.55 | 372 | GLU | 1.46 | 124 | GLU | 1.54 |
| 70 | 230 | GLU | 1.55 | 384 | ASP | 1.44 | 868 | ASP | 1.50 |
| 71 | 393 | MET | 1.54 | 387 | GLU | 1.44 | 869 | ARG | 1.50 |
| 72 | 395 | TYR | 1.51 | 388 | ASN | 1.44 | 400 | VAL | 1.47 |
| 73 | 848 | GLY | 1.48 | 191 | GLU | 1.44 | 1186 | ASP | 1.45 |
| 74 | 849 | THR | 1.48 | 260 | ASP | 1.44 | 1187 | ARG | 1.45 |
| 75 | 1056 | GLU | 1.46 | 1060 | LYS | 1.43 | 934 | LYS | 1.39 |
| 76 | 1060 | LYS | 1.46 | 1219 | ASP | 1.42 | 935 | THR | 1.39 |
| 77 | 776 | ASP | 1.45 | 1220 | ILE | 1.42 | 943 | ARG | 1.39 |
| 78 | 777 | GLY | 1.45 | 873 | GLU | 1.41 | 947 | LYS | 1.39 |
| 79 | 1183 | VAL | 1.41 | 1201 | SER | 1.41 | 1181 | SER | 1.36 |
| 80 | 1184 | ASP | 1.41 | 1202 | ASN | 1.41 | 134 | ASN | 1.34 |
| 81 | 739 | TYR | 1.38 | 1208 | ARG | 1.41 | 260 | ASP | 1.32 |
| 82 | 741 | ASP | 1.38 | 360 | HIS | 1.40 | 261 | TRP | 1.32 |
| 83 | 742 | VAL | 1.38 | 401 | TRP | 1.40 | 262 | GLN | 1.32 |
| 84 | 743 | ASP | 1.38 | 754 | HIS | 1.40 | 263 | GLU | 1.32 |
| 85 | 124 | GLU | 1.38 | 81 | ARG | 1.40 | 264 | GLN | 1.32 |
| 86 | 391 | ASP | 1.36 | 85 | GLU | 1.40 | 1084 | GLU | 1.32 |
| 87 | 394 | GLN | 1.36 | 199 | VAL | 1.40 | 1085 | PRO | 1.32 |
| 88 | 1111 | LEU | 1.36 | 449 | GLN | 1.37 | 1102 | GLN | 1.31 |
| 89 | 152 | GLU | 1.34 | 383 | LYS | 1.33 | 1103 | SER | 1.31 |
| 90 | 263 | GLU | 1.34 | 579 | ARG | 1.30 | 466 | LEU | 1.29 |
| 91 | 831 | TYR | 1.34 | 581 | ASP | 1.30 | 467 | ALA | 1.29 |
| 92 | 579 | ARG | 1.31 | 582 | ASP | 1.30 | 492 | LEU | 1.27 |
| 93 | 580 | LEU | 1.31 | 920 | THR | 1.28 | 933 | SER | 1.27 |
| 94 | 581 | ASP | 1.31 | 924 | ARG | 1.28 | 778 | THR | 1.26 |
| 95 | 489 | GLU | **1.31** | 931 | LEU | 1.26 | 900 | ALA | 1.25 |
| 96 | 491 | ASP | 1.31 | 133 | ASP | 1.26 | 912 | GLY | 1.24 |
| 97 | 1181 | SER | 1.29 | 138 | HIS | 1.26 | 913 | CYS | 1.24 |
| 98 | 536 | GLU | **1.29** | 465 | ASP | 1.24 | 914 | THR | 1.24 |
| 99 | 537 | GLU | **1.29** | 469 | ASP | 1.24 | 268 | GLY | 1.24 |
| 100 | 1185 | ILE | 1.29 | 546 | ARG | 1.23 | 270 | ASN | 1.24 |
| 101 | 1216 | GLU | 1.29 | 134 | ASN | 1.22 | 155 | ASN | 1.20 |
| 102 | 159 | GLN | 1.27 | 137 | GLN | 1.22 | 156 | LEU | 1.20 |
| 103 | 160 | ALA | 1.27 | 468 | ASN | 1.21 | 158 | LEU | 1.20 |
| 104 | 1134 | HIS | 1.26 | 1062 | GLU | 1.20 | 917 | GLY | 1.18 |
| 105 | 535 | GLU | 1.26 | 1184 | ASP | 1.17 | 1196 | ASP | 1.17 |
| 106 | 260 | ASP | 1.25 | 1068 | ASP | 1.15 | 1197 | CYS | 1.17 |
| 107 | 261 | TRP | 1.25 | 1084 | GLU | 1.15 | 1202 | ASN | 1.17 |
| 108 | 1124 | ALA | 1.25 | 1086 | SER | 1.15 | 1206 | MET | 1.17 |
| 109 | 1126 | ASP | 1.25 | 408 | GLN | 1.13 | 1207 | GLU | 1.17 |
| 110 | 1142 | ARG | 1.25 | 885 | THR | 1.09 | 1056 | GLU | 1.15 |
| 111 | 1144 | GLU | 1.25 | 1216 | GLU | 1.09 | 1060 | LYS | 1.15 |
| 112 | 1145 | ASP | 1.25 | 371 | LYS | 1.09 | 952 | GLY | 1.15 |
| 113 | 190 | PRO | 1.22 | 398 | GLN | 1.09 | 953 | ARG | 1.15 |
| 114 | 191 | GLU | 1.22 | 832 | ASP | 1.08 | 248 | GLU | 1.14 |
| 115 | 1221 | TYR | 1.22 | 833 | GLU | 1.08 | 280 | GLU | 1.14 |
| 116 | 148 | LEU | 1.20 | 834 | GLU | 1.08 | 740 | ASN | 1.14 |
| 117 | 1143 | GLU | 1.19 | 1198 | LYS | 1.07 | 741 | ASP | 1.14 |
| 118 | 1146 | ARG | 1.19 | 535 | GLU | 1.06 | 831 | TYR | 1.14 |
| 119 | 133 | ASP | 1.16 | 536 | GLU | **1.06** | 123 | VAL | 1.12 |
| 120 | 137 | GLN | 1.16 | 733 | HIS | 1.05 | 629 | ASP | 1.11 |
| 121 | 396 | CYS | 1.16 | 734 | HIS | 1.05 | 630 | ASN | 1.11 |
| 122 | 1086 | SER | 1.14 | 152 | GLU | 1.03 | 745 | PRO | 1.11 |
| 123 | 192 | GLU | 1.13 | 193 | ARG | 1.03 | 747 | CYS | 1.11 |
| 124 | 348 | LEU | 1.12 | 106 | HIS | 1.01 | 748 | TRP | 1.11 |
| 125 | 351 | SER | 1.12 | 110 | HIS | 1.01 | 830 | ASP | 1.11 |
| 126 | 352 | SER | 1.12 | 1214 | GLN | 1.00 | 773 | LYS | 1.11 |
| 127 | 492 | LEU | 1.12 |  |  | 5.99 | 348 | LEU | 1.10 |
| 128 | 600 | PRO | 1.09 |  |  | 5.99 | 272 | SER | 1.09 |
| 129 | 601 | LYS | 1.09 |  |  | 5.99 | 273 | PHE | 1.09 |
| 130 | 1196 | ASP | 1.07 |  |  | 4.85 | 583 | PRO | 1.09 |
| 131 | 1207 | GLU | 1.07 |  |  | 4.85 | 71 | HIS | 1.07 |
| 132 | 1208 | ARG | 1.07 |  |  | 4.48 | 538 | GLU | 1.06 |
| 133 | 608 | ASP | 1.07 |  |  | 3.68 | 542 | ASP | 1.06 |
| 134 | 950 | ASN | 1.07 |  |  | 3.68 | 344 | SER | 1.06 |
| 135 | 953 | ARG | 1.07 |  |  | 3.68 | 1087 | ALA | 1.06 |
| 136 | 622 | TYR | 1.07 |  |  | 3.40 | 1089 | GLN | 1.06 |
| 137 | 736 | ASN | 1.07 |  |  | 3.37 | 835 | GLY | 1.04 |
| 138 | 369 | LEU | 1.07 |  |  | 3.37 | 540 | GLN | 1.02 |
| 139 | 450 | GLY | 1.03 |  |  | 3.34 | 543 | VAL | 1.02 |
| 140 | 451 | THR | 1.03 |  |  | 3.34 |  |  | 5.14 |
| 141 | 582 | ASP | 1.03 |  |  | 3.22 |  |  | 5.14 |
| 142 | 383 | LYS | 1.02 |  |  | 3.22 |  |  | 4.93 |
| 143 | 384 | ASP | 1.02 |  |  | 2.98 |  |  | 4.93 |
| 144 | 268 | GLY | 1.01 |  |  | 2.98 |  |  | 3.88 |
| 145 | 270 | ASN | 1.01 |  |  | 2.89 |  |  | 3.54 |

**Table S3.**
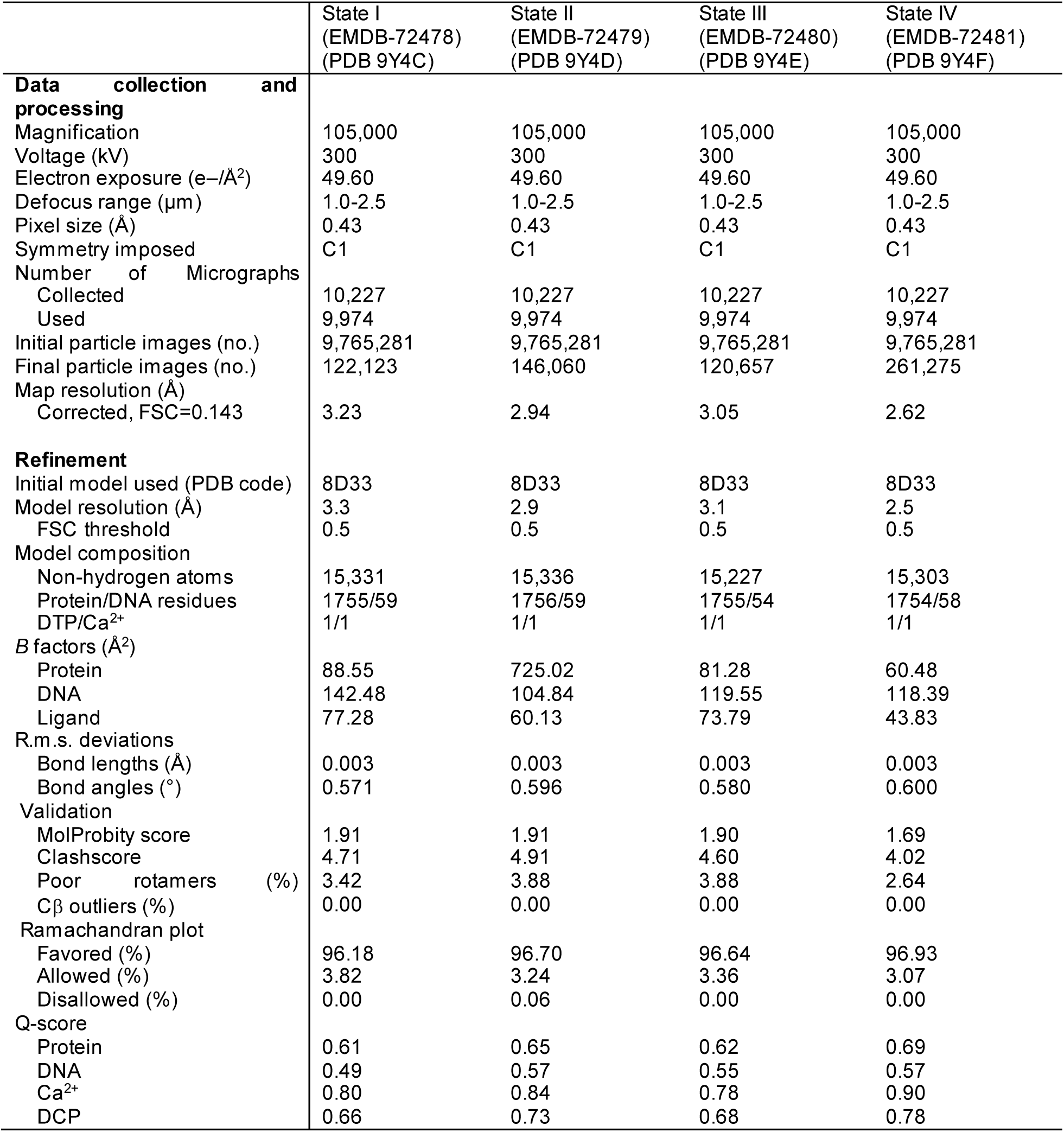
Cryo-EM data collection, refinement, and validation statistics of Pol γ strand-displacement complexes.

**Figure S1.**
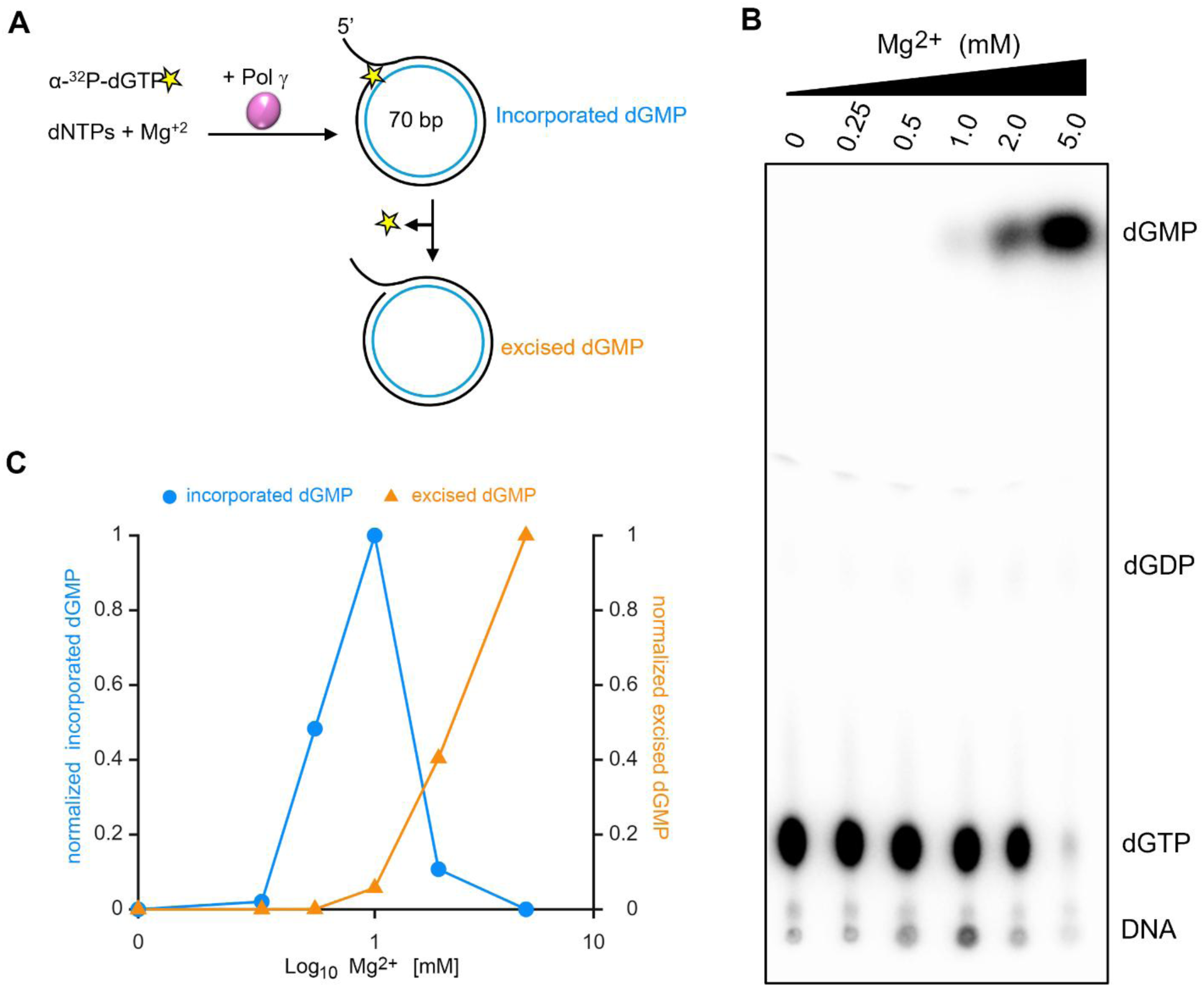
Polymerase and exonuclease activities of Pol γ exo+ measured during ongoing DNA synthesis. (**A**) Schematic of the experimental design used to simultaneously measure polymerase and exonuclease activities of Pol γ during strand displacement rolling circle DNA replication. (**B**) TLC scan shows products of replication reactions performed with stated magnesium concentrations. (**C**) Fractions of dGMP incorporated in the synthesized DNA and fractions of dGMP excised during DNA synthesis reactions performed at different magnesium concentrations. Circle and triangles show mean data from two independent repeats.

**Figure S2.**
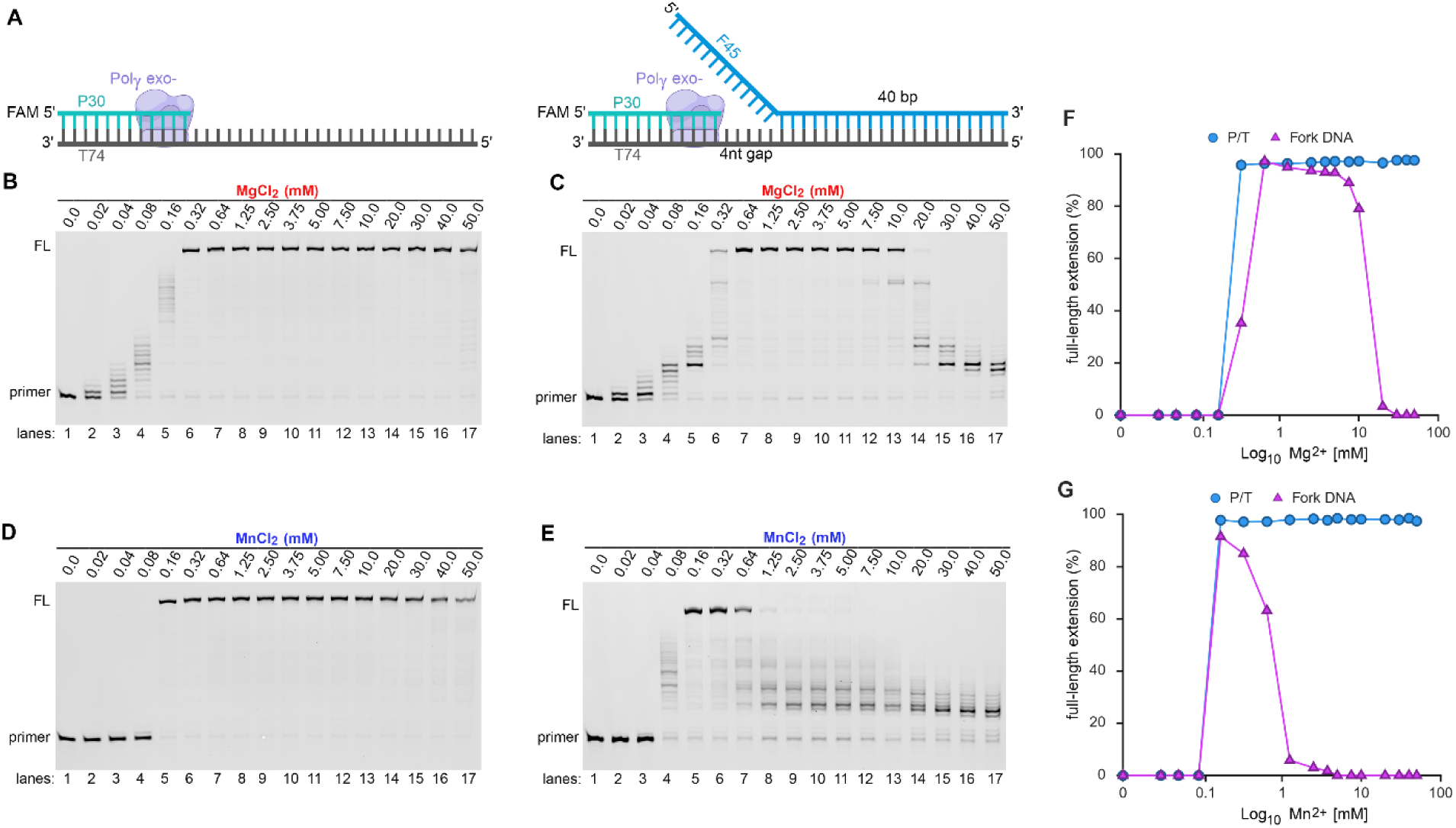
Metal regulation of Pol γ Exo-deficient strand displacement synthesis. (**A**) scheme of the substrate used in this assay. Pol γ Exo-DNA synthesis on p/t (**B**) or fork DNA (**C**) in the presence of 0-50 mM Mg^2+^, or in the presence of 0-50 mM Mn^2+^ (**D** and **E**). Quantification of metal-dependent full-length primer extension in the presence of Mg^2+^ or Mn^2+^ is shown in (**F**) and (**G**).

**Figure S3.**
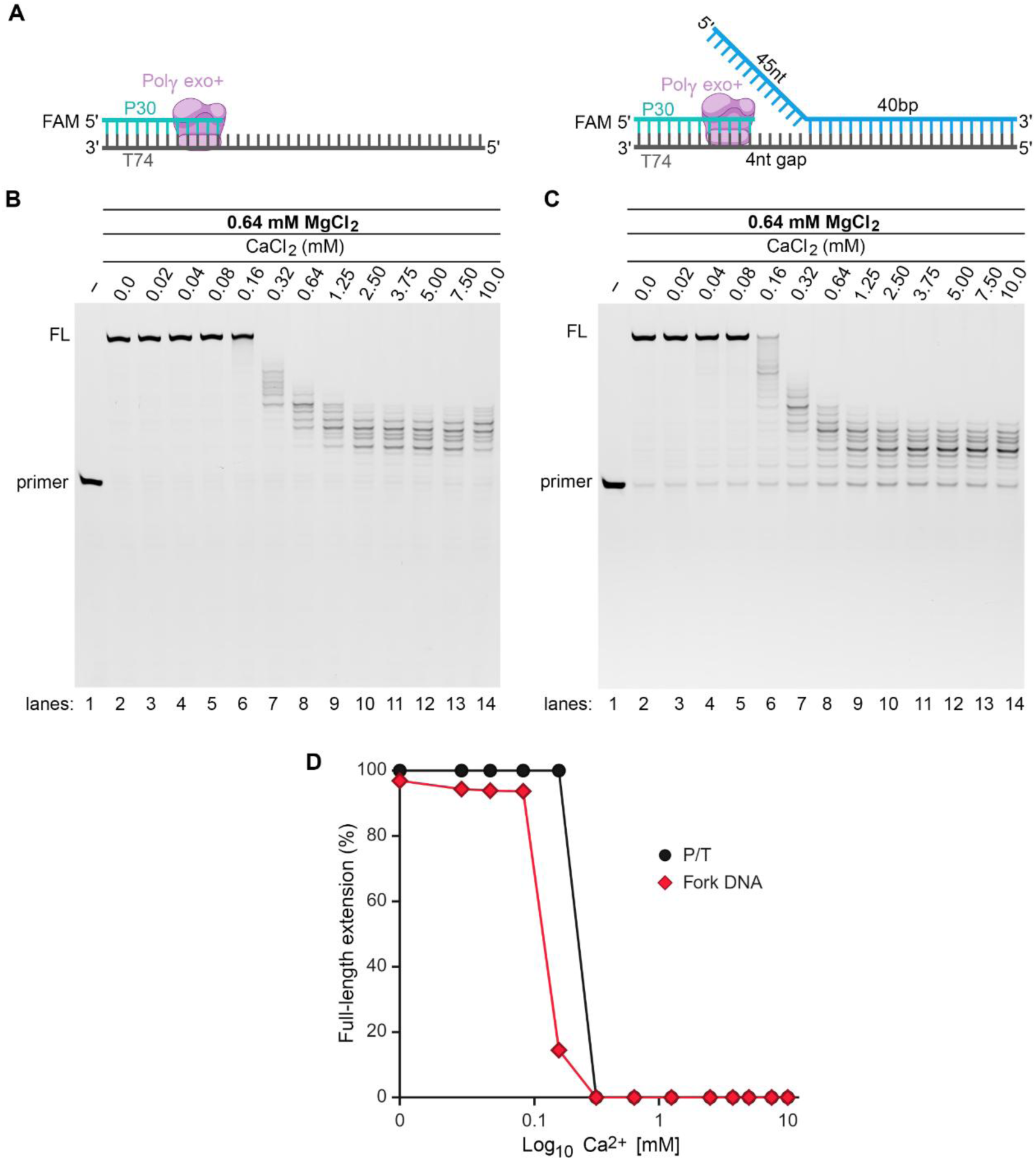
Pol γ strand displacement synthesis in the presence of mixed divalent metal ions. (**A**) p/t and fork DNA substrates. (**B**-**C**) Primer extension of a p/t DNA substrate in the presence of 0 to 10 mM Ca^2+^ without (**B**) and with a constant 0.64 mM Mg^2+^ (**C**), (**D**) quantification of B-C panels. Full extension product in p/t template assays decreased at 0.32 mM Ca^2+^ while the strand displacement synthesis is over 90% abolished when Ca^2+^ concentration reached 0.16 mM.

**Figure S4.**
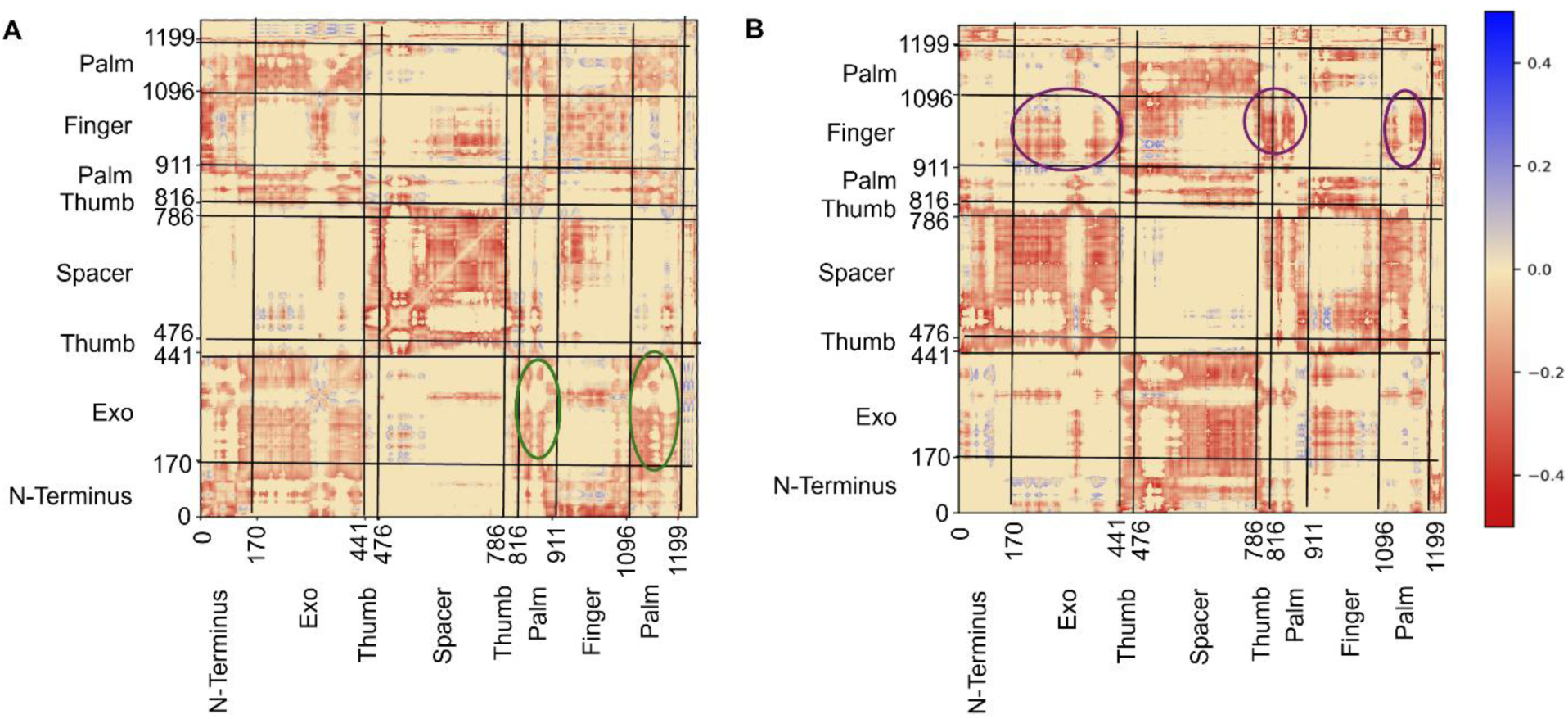
Difference cross-correlation analysis. Correlation (**A**) and anti-correlation (**B**) difference between 20 mM and 1 mM Mg ions. Regions in red show higher correlation/anti-correlation in the 1 mM system with and regions in blue show higher correlation/anticorrelation in the system with 20 mM. The regions with elevated changes are circled.

**Figure S5.**
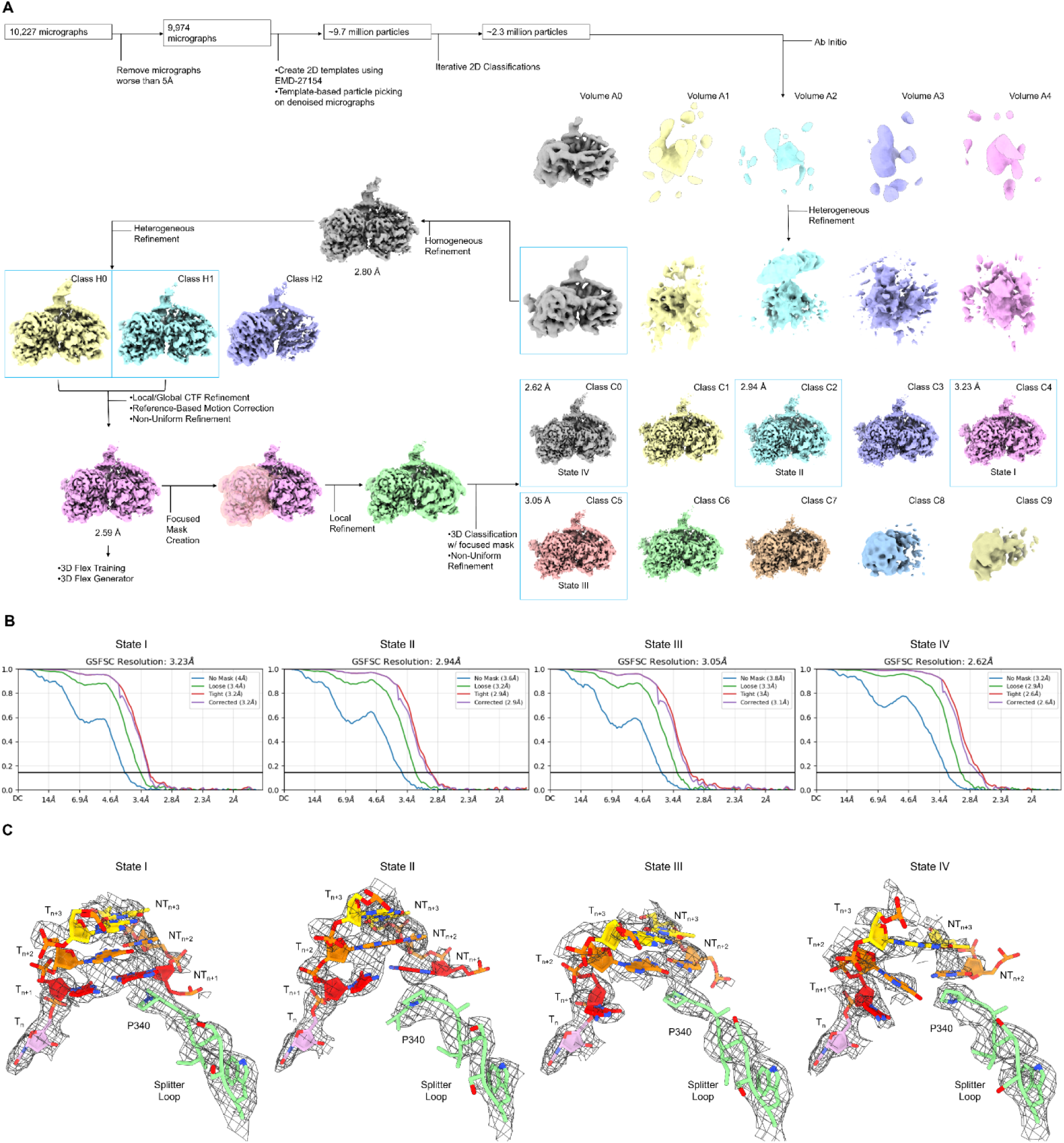
Cryo-EM structural determination. (**A**) Cryo-EM image processing workflow of Pol γ strand displacement complex using cryoSPARC (See Materials and Methods section for details). (**B)** GSFSC resolution curve of each State at 0.143 threshold. (**C**) Electron density map of each state with the corresponding atomic coordinates of the downstream duplex and the Splitter loop.

**Figure S6.**
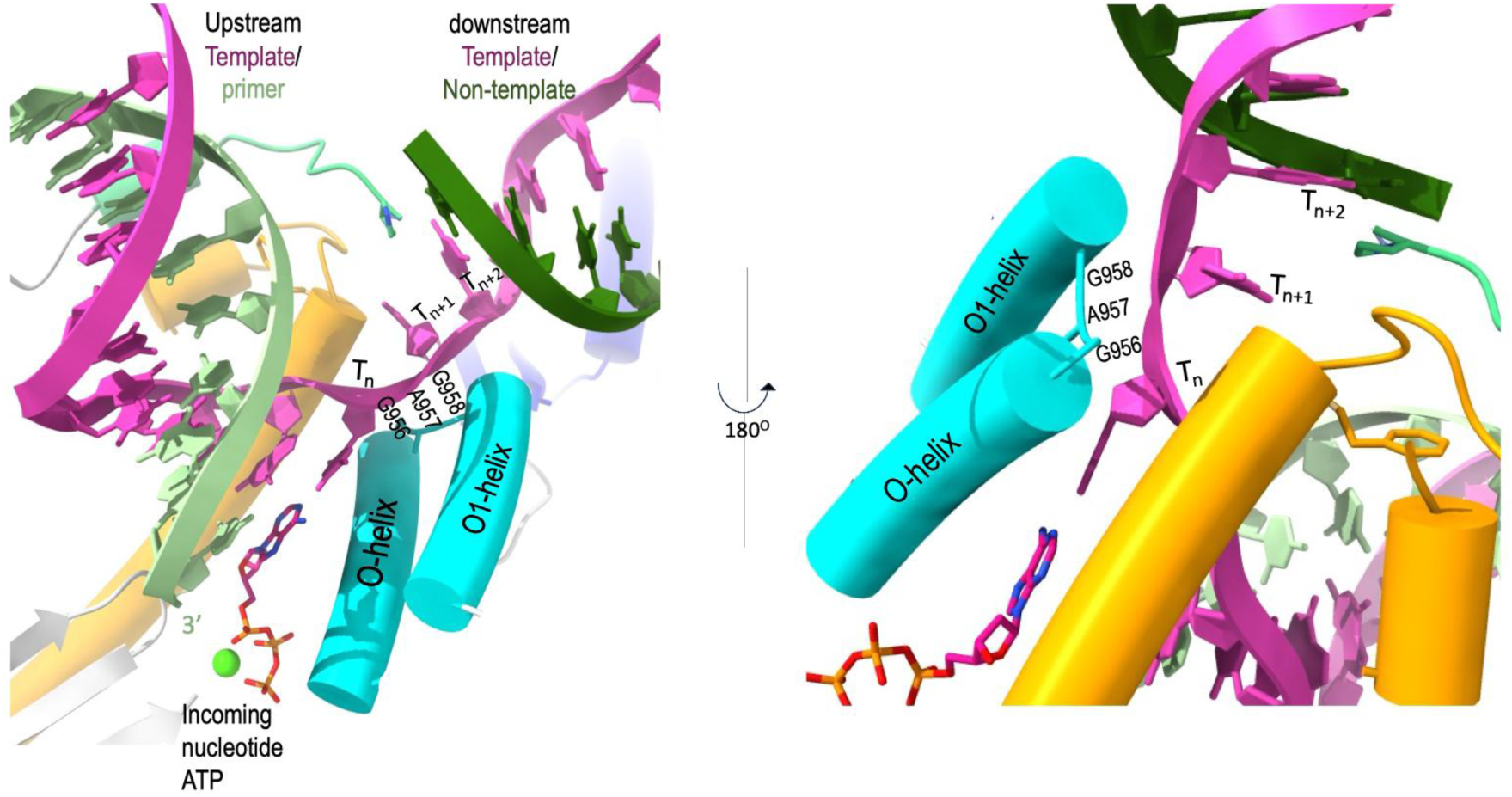
Pol Active Site of Pol γ strand displacement complex. The Pol site of the strand displacement complexes remains unchanged, replication mode with bound an incoming nucleotide forming base pairing interaction with template coding residue Tn. The downstream T_n+1_ and T_n+2_ are stabilized by a loop connecting the O- and O1- helices at residues G^956^A^957^G^958^.

**Figure S7.**
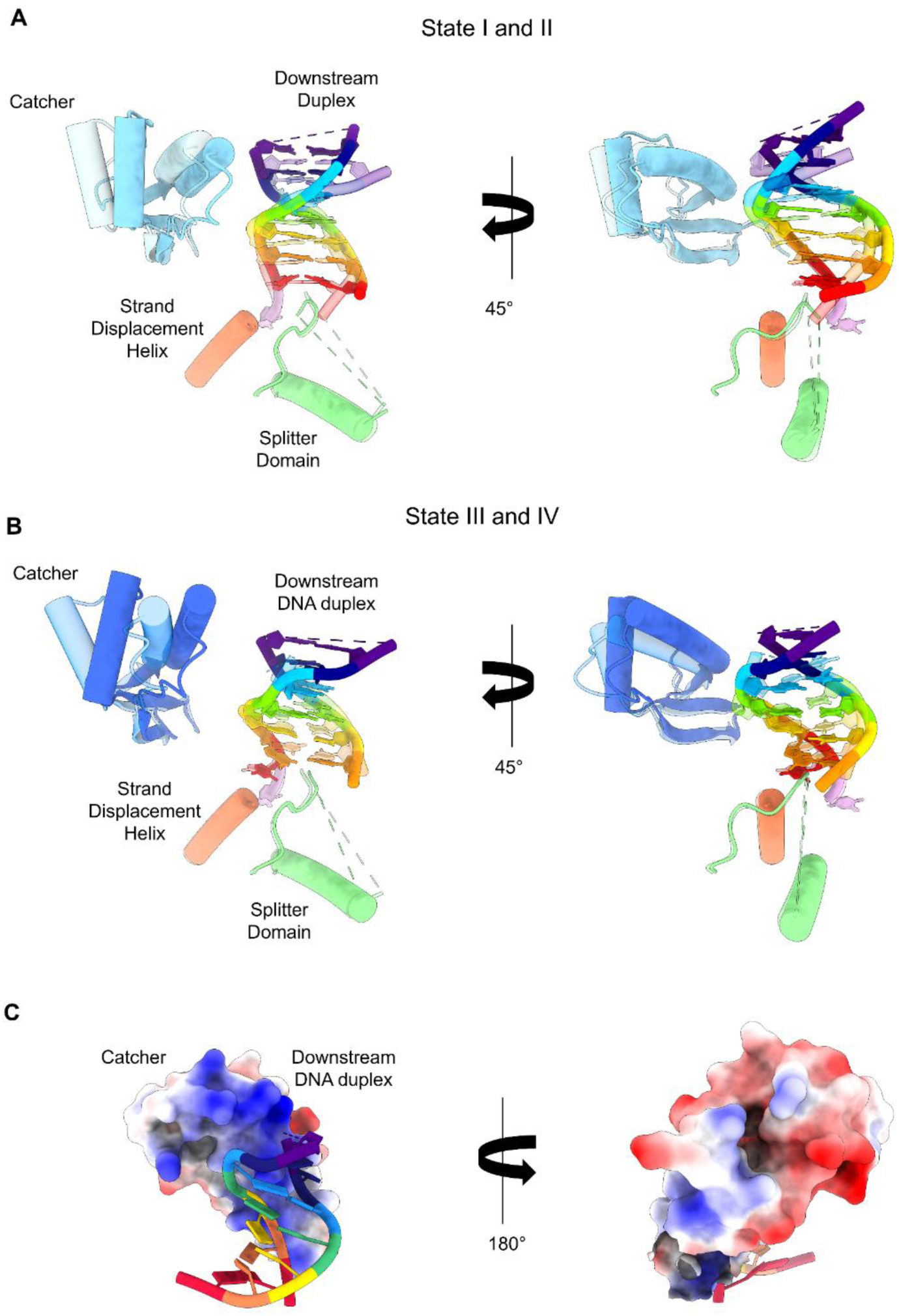
Conformational changes of the Catcher domain among the four States. . Correlated movement of the Catcher domain with the downstream duplex in State I (transparent) and State II (solid) (**A**) and in State III (transparent) to State IV (solid) (**B**). Transitions between state I and II and state III and IV result in scrunching of the downstream DNA duplex that likely facilitates the base pair buckling and eventual unwinding, while allowing Pol γ to remain stationary. (**C**) Electrostatic potential surface of the Catcher domain where the positively charged side (blue) faces the downstream duplex, while the negatively charged area (red) are located on the opposite side.

**Figure S8.**
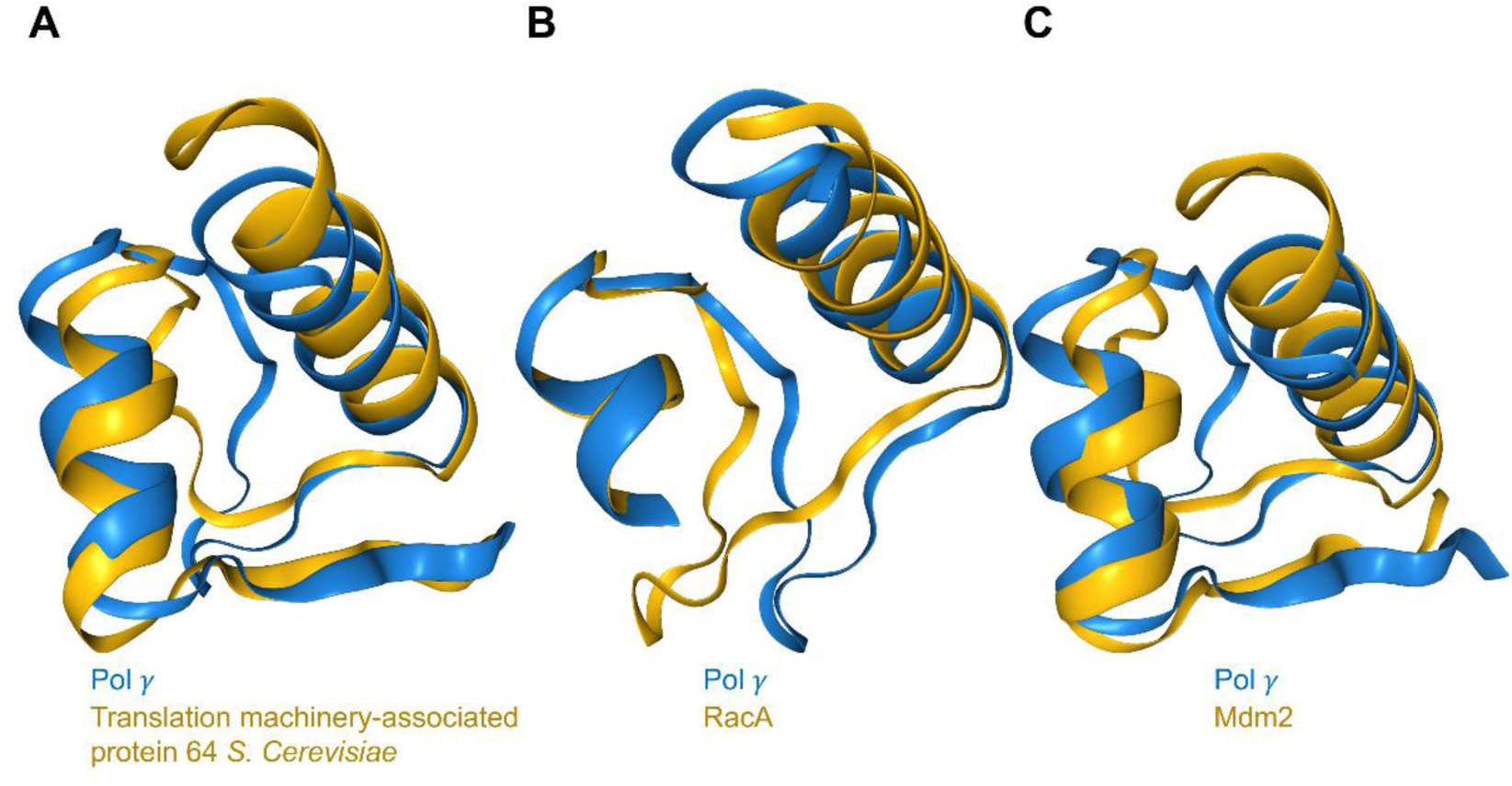
Comparison of the Catcher Domain with Other Proteins with Similar Fold. Pol γ catcher domain (blue) is compared with other proteins that contain similar fold (yellow): Translation machinery-associated protein 64 from *S. Cerevisiae* (**A**), RacA from *B. subtilis* (**B**, PDB: 5I44), and Mdm2 from *H. sapiens* (**C**, PDB: 5ZXF).

**Figure S9.**
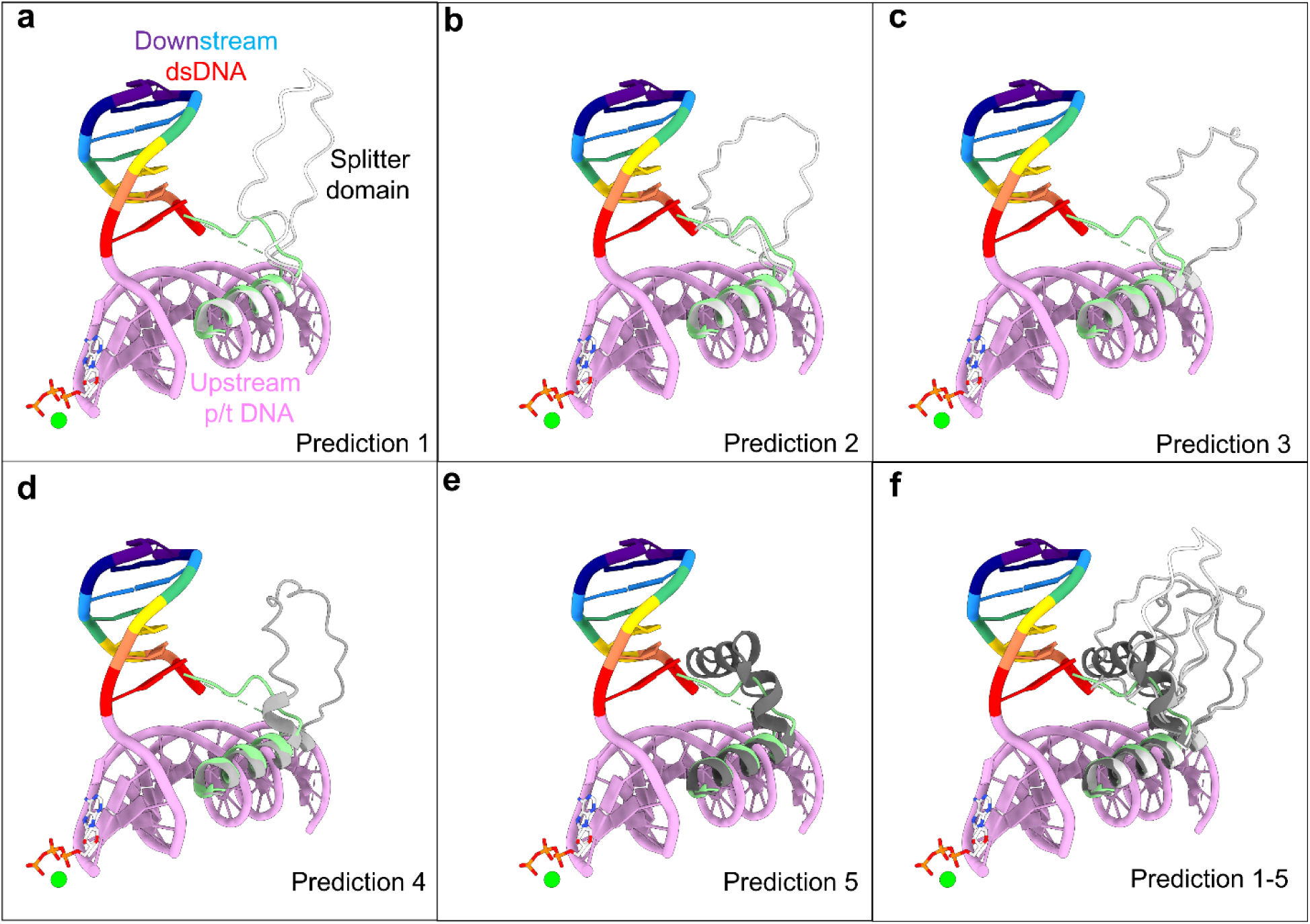
AlphaFold predicted a region of the Splitter domain that was disordered in the cryo-EM structure. Five varied structures were predicted with high to low confidence (Prediction 1 −5). The four high-confidence predictions showed a flexible loop (**a-d**), whereas the lowest confidence prediction showed a helical structure (**e**). **f**. Superposition of all predictions to illustrate the flexibility of the region.

## Notes

### Competing Interest Statement

The authors have declared no competing interest.

